# CRISPRi-based screens in iAssembloids to elucidate neuron-glia interactions

**DOI:** 10.1101/2023.04.26.538498

**Authors:** Emmy Li, Camila Benitez, Steven C. Boggess, Mark Koontz, Indigo V.L. Rose, Delsy Martinez, Nina Draeger, Olivia M. Teter, Avi J. Samelson, Na’im Pierce, Erik M. Ullian, Martin Kampmann

## Abstract

The sheer complexity of the brain has complicated our ability to understand the cellular and molecular mechanisms underlying its function in health and disease. Genome-wide association studies have uncovered genetic variants associated with specific neurological phenotypes and diseases. In addition, single-cell transcriptomics have provided molecular descriptions of specific brain cell types and the changes they undergo during disease. Although these approaches provide a giant leap forward towards understanding how genetic variation can lead to functional changes in the brain, they do not establish molecular mechanisms. To address this need, we developed a 3D co-culture system termed iAssembloids (induced multi-lineage assembloids) that enables the rapid generation of homogenous neuron-glia spheroids. We characterize these iAssembloids with immunohistochemistry and single-cell transcriptomics and combine them with large-scale CRISPRi-based screens. In our first application, we ask how glial and neuronal cells interact to control neuronal death and survival. Our CRISPRi-based screens identified that GSK3β inhibits the protective NRF2-mediated oxidative stress response in the presence of reactive oxygen species elicited by high neuronal activity, which was not previously found in 2D monoculture neuron screens. We also apply the platform to investigate the role of APOE- 4, a risk variant for Alzheimer’s Disease, in its effect on neuronal survival. We find that APOE- 4-expressing astrocytes may promote more neuronal activity as compared to APOE- 3- expressing astrocytes. This platform expands the toolbox for the unbiased identification of mechanisms of cell-cell interactions in brain health and disease.

## Introduction

The human brain contains over 86 billion neurons and approximately the same number of non-neuronal cells^1^, which are all intricately interconnected in the brain. One of the most challenging problems in biology is to disentangle the complexity of the brain and to understand how brain function is altered in disease. Genetic and genomics approaches are uncovering how genetic variation and mutations can drive cellular and molecular changes in the brain. For example, single-cell transcriptomics have revealed the diversity of cell types and cell states in neuronal and glial cells at the molecular level^2^, while genome-wide association studies (GWAS) have identified genetic variants associated with specific neurological phenotypes and diseases^3–5^.

However, these approaches do not elucidate the molecular mechanisms, whether cell autonomous or cell-non autonomous, by which genes control brain function and disease.

To enable the scalable functional characterization of gene function in brain cell types, we previously developed CRISPR-based functional genomics screens in human iPSC-derived neurons^6,7^, microglia^8^,and astrocytes^9^. However, these screens were conducted in monoculture systems and are thus unable to capture mechanisms driven by glia-neuron interactions. Multiple systems have been developed in hiPSC organoid-based culture systems to link specific genetic perturbations to functions^10–12^. A limitation of organoid-based models is the lengthy and labor-intensive process to generate them, and remaining challenges in heterogeneity. To address these challenges, we established a 3D co-culture system we call iAssembloids (induced multi- lineage assembloids) that enables the rapid generation of homogenous neuron-glia spheroids. iAssembloids enable specific control over the origin of cell types of interest, for instance, to combine cells from different genetic backgrounds, enabling the investigation of non-cell autonomous mechanisms. Similar 3D co-culture systems developed previously^13,14^ have been successfully utilized to model features of neurodegenerative diseases such as tau aggregation^15^ and TDP-43 pathology^16^, making our platform expandable to examine other disease-associated paradigms.

Here, we characterize these iAssembloids and combine them with large-scale CRISPRi-based screens to model how glial and neuronal cells can interact to modulate neuronal survival. We also apply the platform to investigate the role of APOE- 4, a major risk variant for Alzheimer’s Disease, in its effect on neuronal survival. Our platform aims to expand our existing functional genomics toolbox to enable a simple, highly scalable method for screening genetic modifiers for neuronal survival in a glial context. Thus, iAssembloids in combination with CRISPRi-based functional genomics will enable the unbiased dissection of mechanisms underlying neuron-glia interactions in health and disease.

## Results

### Neurons, astrocytes, and microglia integrate into three dimensional cultures (iAssembloids)

To generate a scalable three-dimensional neuron-astrocyte-microglia co-culture system that can be used for CRISPRi-based genetic screening approaches, we selected differentiation protocols for hiPSC-derived neurons, astrocytes and microglia that are scalable and rapid. We used the dox-inducible NGN2 system for neuron differentiation^17,18^ and our recently developed six transcription factor (6TF) induction protocol for microglia differentiation^8^. For astrocyte differentiation, we used either of two different protocols, one that is serum-based^19^ and one that is serum-free^13^. For iAssembloid generation, we seeded differentiated astrocytes together with immature differentiated neurons (day 0) at a ratio of 3 neurons to 1 astrocyte into AggreWell^TM^ 800 plates. After one week, we added microglia at a third of the number of astrocytes (Fig. 1A). These cultures self-assemble into spheroids within the wells, are uniform in size (200-400 microns; ∼10,000 cells) and retain their shape over time (Fig. 1B).

**Figure 1.**
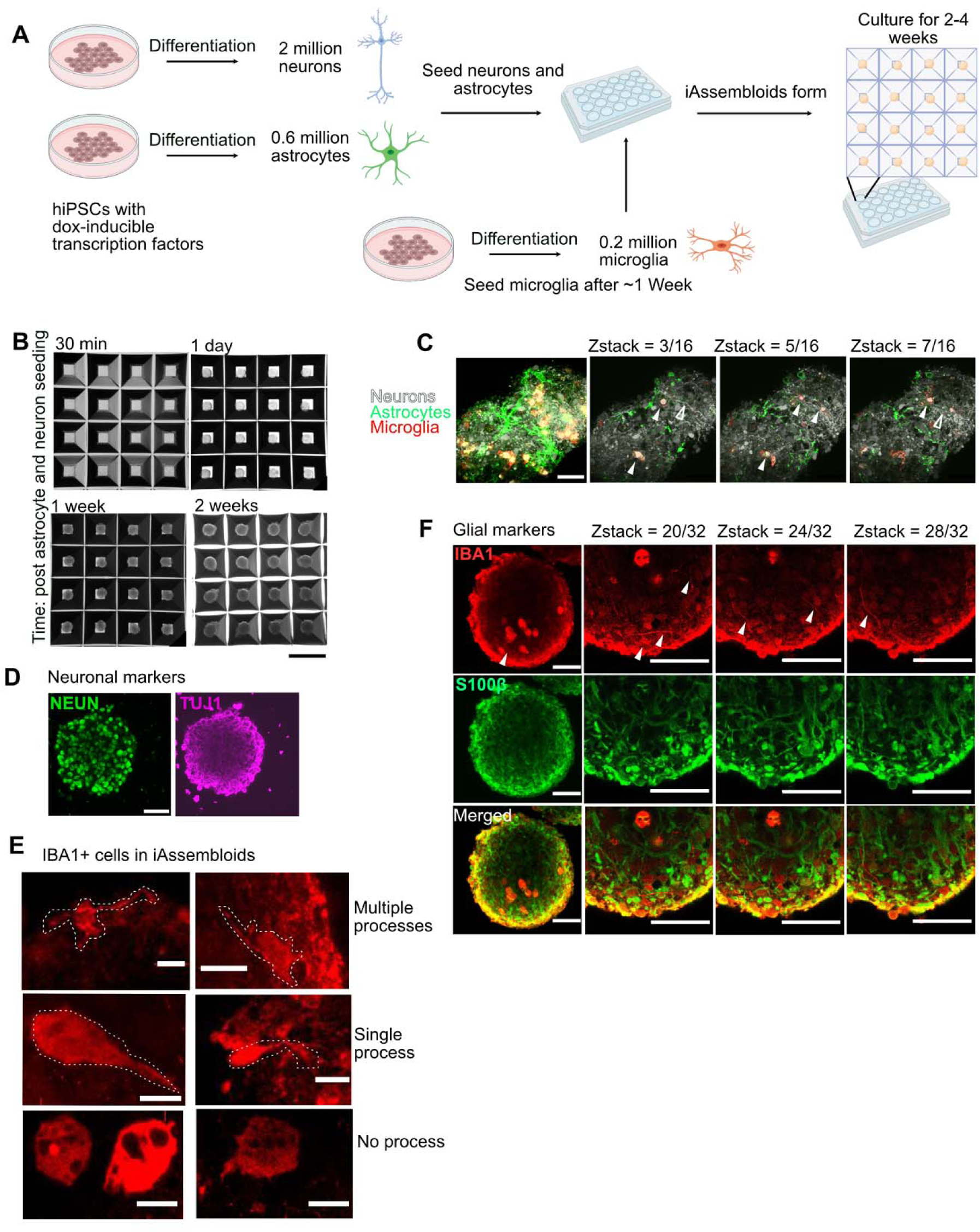
Integrating hiPSC-derived neurons, astrocytes, and microglia into three- dimensional cultures (iAssembloids) **(A)** hiPSCs expressing CRISPRi machinery and inducible NGN2 are pre- differentiated and seeded with hiPSC-derived astrocytes at a 3:1 neurons:astrocyte ratio. After 1 week, hiPSC-derived microglia are seeded at one-third of the number of astrocytes. **(B)** Brightfield images of iAssembloids 30 min, 1 day, 1 week and 2 weeks post seeding. Scale bar = 800 µm. **(C)** Neurons (gray), astrocytes (green) and microglia (red) expressing different fluorescent proteins were seeded to form iAssembloids. Left image: maximum-intensity projection, other images: individual images from the horizontal sample images (z-stack) generated from confocal microscopy. Arrows denote neurons co-localized with microglia (closed arrowheads) as well as neuronal extensions across the culture (open arrowheads). Scale bar = 50 µm. Images were taken 14 days post seeding into AggreWell 800 plates. **(D)** Maximum intensity projections of iAssembloids stained with antibodies against neuronal markers NEUN and TUJ1. Scale bar = 50 µm. Images were taken 14 days post seeding into AggreWell™ 800 plates. **(E)** Maximum intensity projections of iAssembloids stained with antibodies against microglia marker IBA1. Three different morphologies are presented. Scale bar = 10 µm. Cells with processes outlined in white. **(F)** Maximum intensity projections of iAssembloids stained with antibodies against the microglial marker IBA1 and the astrocyte marker S100β. Scale bars = 50 µm. Images were taken 14 days post seeding into AggreWell™ 800 plates. Arrows denote microglia projections.

To visualize the three cell types and their interactions, we seeded iAssembloids (induced multi- lineage assembloids) with hiPSC-derived neurons expressing BFP, astrocytes expressing GFP and microglia expressing mScarlet along with cells expressing no fluorescent proteins. Neurons, astrocytes, and microglia were imaged in the iAssembloids at 14 days post-culture in the AggreWell™ 800 plate. Astrocytes developed a stellate with extensive projections (Fig. 1C). Neurons developed processes that traversed the iAssembloids (Fig. 1C, Video S1). To confirm that all three cell types maintained their cellular identity in iAssembloids throughout differentiation, we stained the iAssembloids for neuronal and glial markers. Neurons within the culture expressed neuronal cell markers NEUN and TUJ1 (Fig. 1D). Microglia, stained with IBA1, fell into two categories: some had few to no processes and migrated towards the middle of the iAssembloid; the others (denoted by outline in E and arrows in F) sent projections through the iAssembloid (Fig. 1E-F). Astrocytes stained positively for S100β (Fig. 1F). Taken together, these data show that the three cell types integrated into the iAssembloids developed complex neuronal and glia morphologies.

### Neurons cultured in iAssembloids are more active than neurons in 2D monoculture

We performed single-nucleus RNA sequencing to further characterize each cell type in the iAssembloids. Nuclei were extracted from whole iAssembloids prior to single-nucleus RNA sequencing. To account for background RNA levels, droplets with less than 500 UMIs were removed. Clusters identified were based on expression of cell type-specific markers (Fig. S1A- B). Cells within iAssembloids expressed appropriate cell type-specific markers (neurons: *RBFOX3, MAPT*; astrocytes: *SLC1A3*; microglia: *SPI1, CSF1R*) and maintained their identity throughout differentiation (Fig. 2A). We found that neurons mainly expressed markers for excitatory neurons (*SLC17A7, SLC17A6*), and not for inhibitory neurons (*GAD1, GAD2, SST, VIP*) (Fig. S1C). We then compared gene expression profiles of neurons cultured within iAssembloids to monocultured neurons and neurons co-cultured with astrocytes that were harvested at the same time point (day 14 post-differentiation)^20^. We computationally mapped neurons from iAssembloids and 2D neurons to a reference dataset of excitatory neurons from human fetal brain tissue^21^. Neurons in iAssembloids mapped most closely to neurons from later gestational weeks (100% to gestational week 17-18) whereas in 2D monoculture neurons, neurons showed a slight shift towards earlier gestational stages (35% to 18 weeks, 57% to 17 weeks, 6% to 13.3 weeks, and 2% to 11.5 weeks (Figure 2B). Using GO term enrichment analysis, we found that genes related to axon guidance and synaptic transmission were more highly expressed in neurons cultured in iAssembloids compared to monoculture (Fig. S1C, Table S1). More specifically, glutamate receptor subunits such as *GRIA1* and *GRIA2* as well as ion-channel subunits (*CACNA1C, KCNQ1, KCNIP4*) were more highly expressed in neurons cultured in iAssembloids (Fig. 2C, Fig. S1D, Table S1). Similarly, we compared neurons from our culture system to a previously published system where neurons were co-cultured with astrocytes for 14 days and also found higher expression of glutamate receptor subunits (Fig S1E-F). Therefore, we hypothesized that neurons cultured in iAssembloids may have higher levels of neuronal activity than neurons in monoculture. We first assayed whether neurons in iAssembloids had higher levels of c-FOS, an immediate early gene whose expression is triggered by neuronal activity, Indeed, we saw that there was a higher level of c-FOS in neurons with iAssembloids as compared to 2D monoculture neurons (Fig. S1G). We therefore assessed the neurons’ electrophysiological activity by performing multi-electrode array analysis in the two culture conditions (Fig. 2D). After two weeks in culture, neurons in iAssembloids had tenfold greater number of spikes on average. In addition, we find that neurons cultured in 3D by themselves have the highest number of spikes. The number of spikes decrease with the addition of more astrocytes, suggesting that astrocytes play an important role in buffering neuronal hyperexcitability (Fig. 2E). Taken together, these results suggest that neurons cultured in iAssembloids are more electrophysiologically active than those cultured in monoculture.

**Figure 2.**
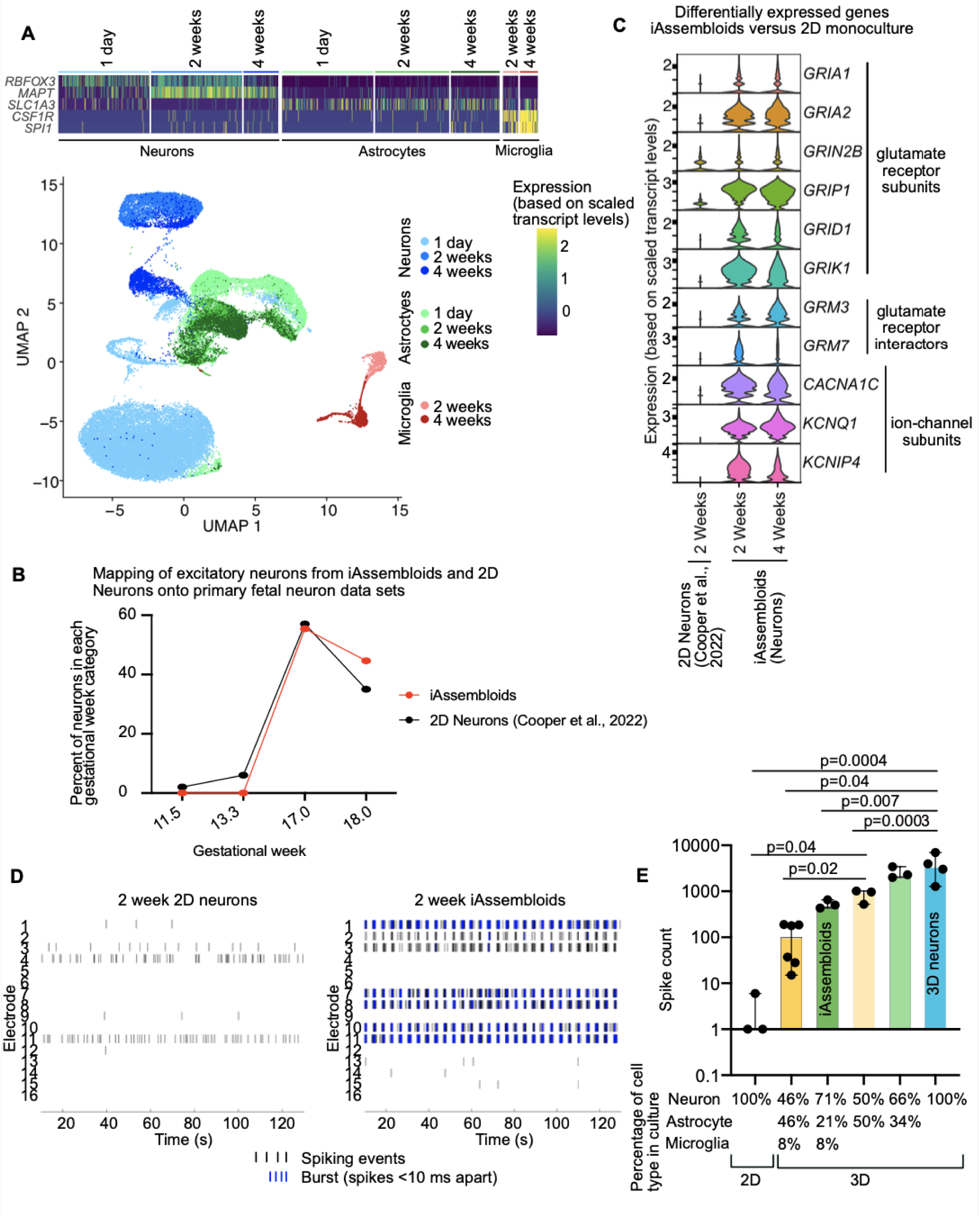
Neurons cultured in iAssembloids are more functionally mature than monocultured neurons **(A)** Single-nucleus RNA sequencing was performed on iAssembloids in culture for 1 day, 2 weeks and 4 weeks. In total, 43,182 nuclei passed quality control and are represented in this dataset. Cell types were assigned to clusters using cell type-specific markers such as *RBFOX3* and *MAPT* for neurons, *SLC1A3* (GLAST) for Astrocytes, and *CSF1R* and *SPI1* for microglia. The heatmap is subsampled at 5,000 cells for visualization. UMAP is labeled by cell type and sample. **(B)** Single-nucleus RNA sequencing of neurons in iAssembloids and 2D monoculture neurons were mapped onto excitatory neurons from fetal brain single-cell sequencing data^21^. The proportion of cells that mapped onto represented gestational weeks are reported. **(C)** Analysis of selected differentially expressed genes comparing iAssembloid and monocultured neuronal culture systems shows that expression of glutamate receptor subunits, glutamate receptor interactors and various ion channel subunits is significantly higher in iAssembloids versus monoculture^20^. **(D)** Example raster plot from multi-electrode array (MEA, Axion) analysis of spikes of neuronal activity in monocultured neurons versus iAssembloids. **(E)** Bar graphs represent cumulative spike data over a 15-minute time span from 2D monoculture and 3D cultures with various proportions of glia. Bars represent the median and the error bars represent the 95% confidence interval. Each dot represents a separate well in the MEA plate.

### Astrocytes in iAssembloids express genes important for supporting neuronal function

Compared to 2D monocultured astrocytes previously generated in our lab^9^, astrocytes cultured in iAssembloids developed distinct transcriptional profiles (Fig. S2A). The clusters that overlapped between the two conditions were either in *TOP2A*+ proliferative clusters or clusters without definitive markers. As astrocytes within iAssembloids matured, astrocytes had transcriptional profiles more strongly associated with supporting neuronal function based on EnrichR pathway and gene ontology analysis of the upregulated genes (Fig. S2B). These functions include BDNF signaling (*JUN, BTG1, PEG10, TUBB*), axon guidance (*ROBO2, NRXN1, NCAM1*), as well as synaptic transmission (*APP, GRID2, NRG3*) (Fig. S2B-C, Table S1). Single-nucleus RNA sequencing also revealed that some astrocytes expressed higher levels of genes associated with glycolysis (*PFKFB4, PDK1, ENO1*) as well as the unfolded protein response (*HSPA5, DDIT3, HERPUD1*) (Fig. S2B-C, Table S1). Very few (0.4-0.5%) of the iAssembloid-derived astrocytes expressed markers of reactive astrocytes such as *IL6*, *IL1B*, *VCAM1*, *CXCL2*, or *CXCL10*^9,23^, suggesting that astrocytes in iAssembloids do not spontaneously become inflammatory (Fig. S2D, Table S1). We validated this by comparing GFP+ astrocytes in iAssembloids with and without the microglial cytokines IL1a+TNF+C1q. Imaging showed a lower number of processes between astrocytes induced by IL1a+TNF+C1q as compared to those found without stimulation (Fig S2E). These results suggest that astrocytes develop homeostatic signatures in iAssembloids that are not represented in monoculture.

### Microglia populations unique to iAssembloids express major histocompatibility complex genes

To characterize transcriptional changes that occur in microglia cultured in iAssembloids, we compared single-nucleus RNA-sequencing data for microglia in iAssembloids to those for monocultured iPSC-derived microglia^8^ (Fig. S3A). We found that 95% of the microglia in day 14 iAssembloids (iMicroglia age 13 days, 5 days in iAssembloid culture) fell into clusters also seen in monocultured microglia. By 28 days in iAssembloid culture, 79% of microglia fell into unique clusters. These clusters, which we named “iAssembloid *SPP1*+”, “iAssembloid chemokine”, and “iAssembloid proliferative”, have profiles like the monocultured microglia *SPP1+*, chemokine, and proliferative clusters, but are characterized by expression of MHC class II proteins such as *HLA-DRA* and *HLA-DRB1* (Fig. S3B, Table S1), which were either not expressed or not captured by microglia in the monoculture dataset. Components of MHC class II pathway have been shown to be expressed by microglia in the context of the brain in healthy human brains, and increase in expression in diseases such as Alzheimer’s Disease or Multiple Sclerosis^24,25^. We mapped our dataset onto an existing human patient dataset^26^ and found that expression of *HLA-DRA* and *HLA-DRB1* between microglia within the iAssembloids and patient microglia were comparable (Fig. S3C). Therefore, iAssembloids provide a model system to study this disease-relevant state of human microglia.

### CRISPRi-based screens identify iAssembloid-specific factors in neuronal survival

As neuronal death is a key hallmark of neurodegenerative disease, we wanted to probe which pathways can make neurons vulnerable and which can protect neurons from neuronal death. While we previously conducted CRISPRi-based modifier screens for neuronal survival in 2D monocultured neurons^6,7^, the more physiological context of our iAssembloid model motivated us to conduct similar screens in iAssembloids. To perform the screen, hiPSCs expressing the dox-inducible transcription factor, NGN2, and dCas9-KRAB CRISPRi machinery were transduced with 19,000 sgRNA targeting over 2,900 genes, including kinases and druggable targets (H1 library)^27^, as well as neurodegeneration-related genes from GWAS (neurodegeneration library, Table S2). To determine genes essential for neuronal survival, we induced neuronal differentiation, incorporated these cells into iAssembloids and compared sgRNA frequencies in the starting population (1 day in culture) and later timepoints (14 and 28 days in culture) to uncover genes controlling neuronal survival (Fig. 3A, Table S3). Many of the hits fell into categories related to metabolism, neuronal activity, cell adhesion, lipid and lipoprotein function, immune response, stress response, and endocytosis (Fig. 3B). Functional categories of hits were identified using the DAVID GO platform, with all targeted genes as the background list and genes determined as hits as the test gene list. We used DAVID’s functional clustering platform and assigned shortened names to the clusters, which are defined as the categories, and used the enrichment scores for the top categories for annotation (Fig. S4A, Table S3). We confirmed that our screen was highly reproducible across two different time points (Fig. S4B). We also performed the screen using two different astrocyte protocols to demonstrate that our iAssembloid system performs comparably with either of the protocols. One astrocyte differentiation protocol takes 6 months to generate the astrocytes, while the other protocol takes only 30 days, which has a practical advantage. CRISPRi screens using the shorter protocol for astrocytes yielded similar results to screens using the more laborious protocol (Fig. S4C).

**Figure 3.**
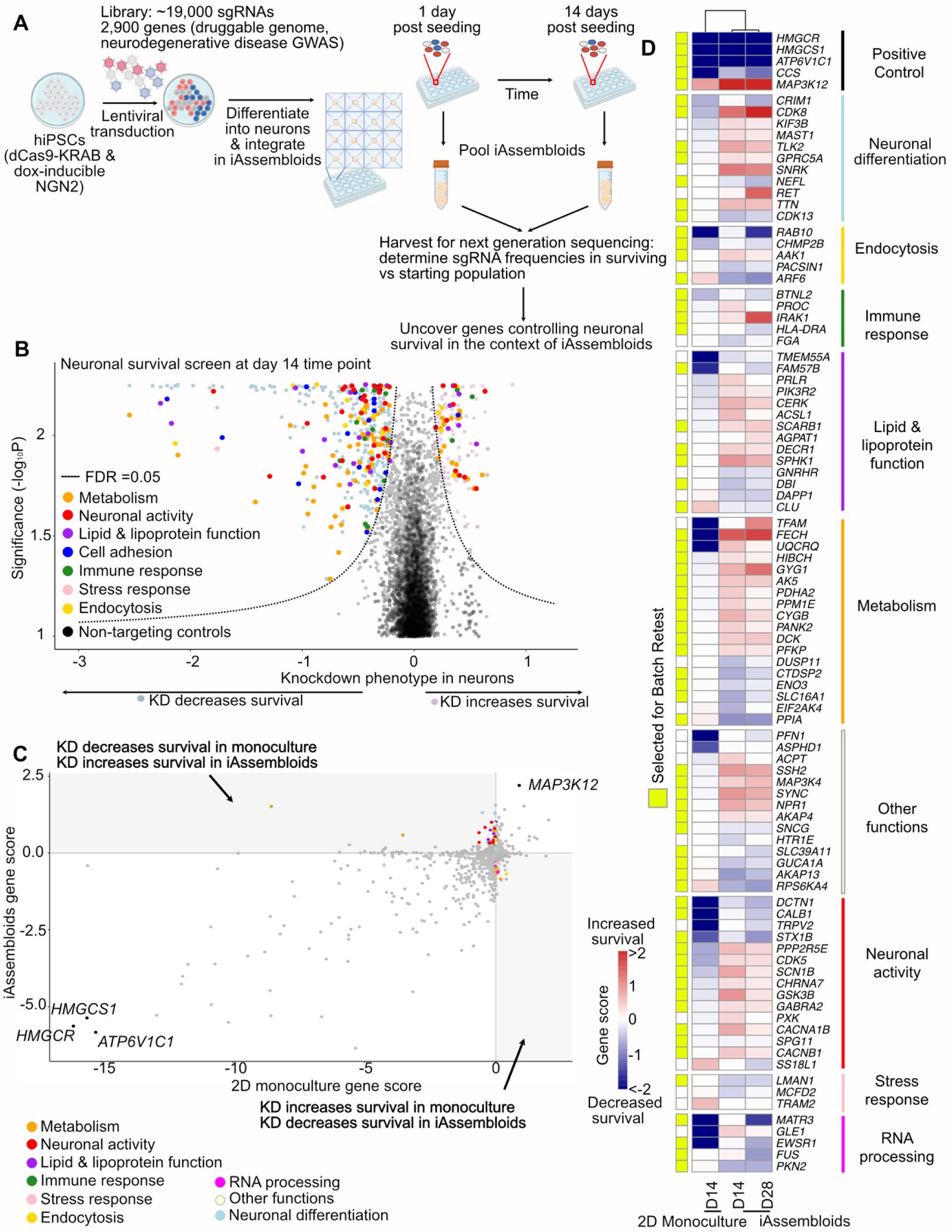
CRISPRi-based screens in iAssembloids reveals genes affecting neuronal survival not found in screens in 2D monoculture. **(A)** Schematic of screen design. **(B)** Volcano plot of knockdown phenotypes from the survival screen from day 14 iAssembloids. Hit genes (FDR < 0.05) are labeled in either light blue (knockdown decreases survival) or light pink (knockdown increases survival). Selected hit genes were color-coded based on curated functional categories. **(C)** Scatterplot of gene scores from our previous survival screen in 2D monocultured neurons^6^ (x-axis) compared to the screen in iAssembloid (y-axis). Both screens used the H1 sgRNA library targeting the “druggable genome.” Selected genes with consistent phenotypes in both screens are labeled in black. Other genes of interest are color-coded based on functional category as in panel B. **(D)** Heatmap representing gene scores for neuronal survival from screens in 2D monoculture screens^6,7^, and from iAssembloids (this study, screens were conducted with the H1 sgRNA library and an sgRNA library targeting neurodegeneration-related genes). Neurodegeneration library hits were compared to gene scores from a genome-wide 2D monoculture screen^7^. Genes are grouped by functional categories as in panels B and C. Genes selected for secondary screens are highlighted by yellow boxes on the left.

We compared our findings to those of screens previously performed in neuron monocultures in our lab^6,7^. The screen using the H1 library were compared to Tian et al., 2019, while the screen using the neurodegeneration library was compared to the genome-wide screen from Tian et al., 2021. Based on Pearson correlation of gene scores from survival screens of 2D neurons versus iAssembloids, iAssembloid screens conducted at two different timepoints were highly correlated with each other, but not as highly correlated with the original 2D neuron screen (Fig. S4D). Essential genes for neuronal survival identified in monoculture, such as *HMGCR*, *HMGCS1* and *ATP6V1C1,* were also found in our iAssembloid screens (Fig. 3C). We selected a subset of hit genes of interest that were iAssembloid-specific and had the same phenotype at day 14 and day 28 in the iAssembloids (Fig. 3D). These hits represent knockdowns that had opposing phenotypes in monoculture versus iAssembloid screens, taking into consideration both the fold change and the significance of the hit, as well reproducibility across the two time points. In total, we curated a list of 68 genes that fit the criteria for further validation (Table S3).

### Focused validation screens assess effect of cell type and media composition on neuronal survival in iAssembloids

We conducted a focused validation screen targeting the 68 curated genes from the primary screen. Comparing the focused validation screen conducted with the same conditions as the primary screen, we found that the results were highly correlated (r=0.9) (Fig. S5A, Table S3). As media composition is altered when microglia are seeded, we assessed whether changing media composition affected neuronal survival in the iAssembloids. In addition, we tested BrainPhys media as it has been shown to increase neural electrical activity in culture^28^. Through hierarchical clustering, we found that screens conducted with differing media compositions were more closely related to each other, except for the screen conducted in BrainPhys media. Thus, except for BrainPhys media, there did not seem to be a strong influence on survival phenotype based on media. We also compared iAssembloids with neurons co-cultured with microglia and 3D neuronal monocultures and found that with 3D neuron monoculture clustered more closely to the screen conducted in BrainPhys media (Fig. S5B, Table S3). To understand the downstream effects on neuronal function after perturbation of these genes, we selected hits that maintained a phenotypic difference between monoculture and iAssembloid culture for a focused screen with single-cell transcriptomic readout (Fig. S5B).

### CROP-seq (CRISPR droplet sequencing) suggests that GSK3 (Glycogen Synthase Kinase 3 Beta) influences neuronal NRF2 activity in the iAssembloid system

To assess transcriptomic changes resulting from genetic perturbations and to uncover pathways that are differentially regulated between iAssembloids and monocultures, we performed CROP-seq (CRISPR droplet sequencing)^29^. After transduction with a pooled sgRNA library and neuronal differentiation, we enriched the neuronal population using magnetic activated cell sorting (MACS) and then we performed single cell sequencing on the enriched neuronal population (Fig. 4A, Table S4) from iAssembloids and 2D monoculture.

**Figure 4.**
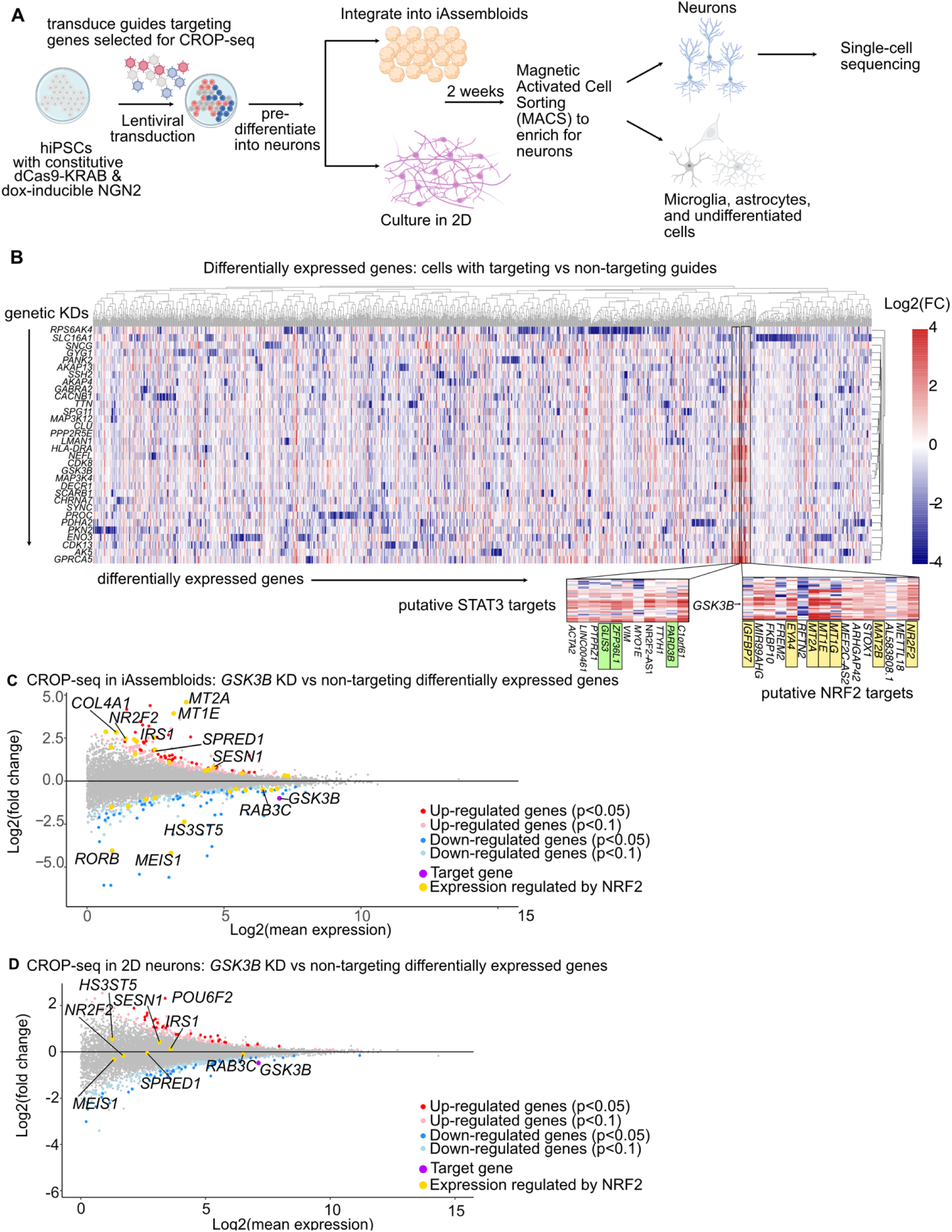
CROP-seq screen reveals that *GSK3B* knockdown induces neuronal expression of NRF2 target genes in iAssembloids, but not 2D monoculture **(A)** Schematic of CROP-seq experimental design. **(B)** Heatmap of combined significantly expressed genes (p<0.05) from CROP-seq in neurons cultured in iAssembloids. Rows represent genes that were knocked down in the cells, columns are differentially expressed genes in cells with knockdowns compared to controls. Highlighted are upregulated genes that are putative NRF2 targets (yellow) and STAT3 targets (green). **(C)** Differentially expressed genes comparing *GSK3B* knockdown versus cells containing non-targeting controls in iAssembloids were determined. Red dots represent upregulated genes below the p<0.05 cutoff whereas pink dots represent genes below the p<0.1 cutoff. Dark blue represents downregulated genes meeting p<0.05 cutoff whereas light blue represents downregulated genes below the p<0.1 cutoff. The target gene, *GSK3B*, is highlighted in purple and genes that have been putatively shown to be regulated by NRF2 are highlighted in yellow. P values were determined using the Wald test. **(D)** Differentially expressed genes comparing *GSK3B* knockdown versus cells containing non-targeting controls in 2D neuronal monoculture were determined as specified in C.

We found that knockdown of several genes, including *GSK3B, TTN, LMAN1* and players in the JNK MAPK pathway drove up expression of different targets of the transcription factors NRF2 (Nuclear factor erythroid 2-related factor 2) and STAT3 (Fig. 4B). We were uniquely interested in the relationship between GSK3 and NRF2 because GSK3 is hypothesized to be highly relevant in a variety of neurodegenerative diseases^30,31^. GSK3 is a kinase that has been proposed to regulate many different cellular functions, ranging from cellular differentiation to apoptosis^32^. In Alzheimer’s disease, especially, GSK3 has been shown to phosphorylate the protein tau^33,34^. Phosphorylation of Tau has been suggested to drive Tau aggregation, leading to neuronal death and the progression of disease^35^ while NRF2 regulates genes in response to oxidative stress^36^. When comparing the effect of *GSK3B* knockdown on neurons in iAssembloids (Fig. 4C, Table S4) and in 2D monoculture (Fig. 4D, Table S4), we found differential regulation of NRF2 targets such as some metallothioneins^37,38^ such as *MT2A, MT1E,* and other genes identified through ChEA Transcription Factor Targets^39^ as NRF2 targets, such as *RORB, HS3ST5,* and *MEIS1,* only in iAssembloids. We therefore concluded that GSK3 may influence NRF2 activity selectively in the iAssembloid system.

### Knockdown of *GSK3B* promotes NRF2 activity by promoting NRF2 nuclear localization

To investigate the mechanism underlying the effect of *GSK3B* knockdown on NRF2, we first confirmed that GSK3 protein levels were reduced by the sgRNAs used in the CROP-seq screen in iAssembloids (Fig. 5A). We next validated the differential effect of *GSK3B* knockdown on neuronal survival in different contexts. We co-cultured cells expressing non-targeting control sgRNAs (GFP+) with cells expressing sgRNAs targeting *GSK3B* (BFP+) at equal ratios and examined changes in those ratios at pre-differentiation and 14 days post-culture. Strikingly, neuronal survival in iAssembloids and in 3D neuronal culture is greatly improved after *GSK3B* knockdown (Fig. 5B), whereas there was no effect of *GSK3B* KD in 2D-monocultured neurons. Thus, the beneficial effect of *GSK3B* knockdown was dependent on 3D culture, but not on the presence of glia.

**Figure 5.**
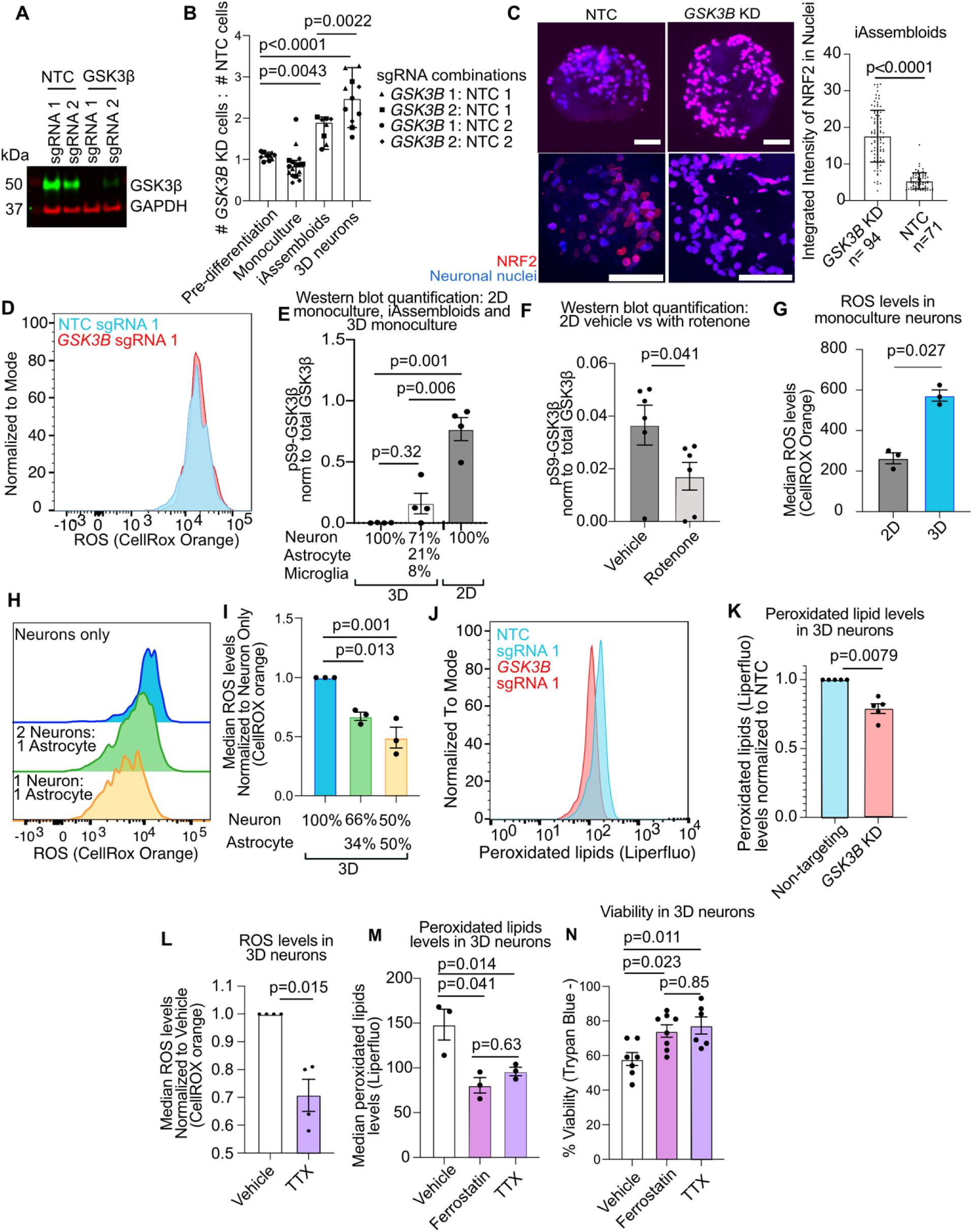
GSK3 activity prevents NRF2 from protecting neurons from oxidative stress **(A)** Western blot validation of GSK3 knockdown in 3D Neurons. NTC: non-targeting control sgRNA. 3D cultures generated from NTC cells from results for 2 independent non-targeting control (NTC) sgRNAs and 2 independent sgRNAs targeting *GSK3B* are shown. **(B)** Validation of the effect of *GSK3*Β knockdown on cell survival over 14 days comparing the ratio of surviving cells expressing a non-targeting control sgRNA and GFP versus the proportion of cells expressing an sgRNA targeting *GSK3*Β and BFP. Experiment was performed with two different NTC sgRNAs as well as two *GSK3*Β sgRNAs with a total of four different combinations of NTC and *GSK3*Β sgRNAs. Medians and 95% confidence intervals are represented. ANOVA followed by Tukey’s multiple comparison test was conducted to determine significance. **(C)** Immunohistochemistry staining for NRF2 (red) in iAssembloids. Neuronal nuclei express BFP (blue). Scale bars = 50 µm. Images taken at 14 days post seeding of iAssembloids into AggreWell^TM^ 800 plates. Nuclei were identified using BFP in images and the integrated intensity for NRF2 within nuclei was quantified. Images from three different iAssembloids expressing NTC sgRNA 1 (n = 71 cells) and *GSK3*Β KD sgRNA 1 (n = 94 cells) were taken. Mean with standard deviation is represented. P values were determined using Student’s t-test. **(D)** Levels of reactive oxygen species (ROS, via CellROX^TM^ orange staining) determined via flow cytometry of neurons expressing a non-targeting control sgRNA or an sgRNA knocking down *GSK3*Β in 3D monoculture. NTC sgRNA 1 and *GSK3*Β KD sgRNA 1 were used in this experiment. Neurons with those sgRNAs were co-cultured and sgRNA identity was assigned based on expression of a BFP (*GSK3*Β sgRNA 1) or mClover (NTC sgRNA 1) marker. **(E)** Western blot quantification comparing level of phospho-S9 GSK3 to total GSK3 in 3D monoculture neurons only, iAssembloids, and 2D monoculture neurons. N=4 independent wells. bars represent mean, error bars represent standard error of the mean. ANOVA followed by Tukey’s multiple comparison test was conducted to determine significance. **(F)** Western blot quantification of pS9-GSK3 levels in 2D monocultured neurons after induction of oxidative stress with rotenone (200 nM for 24 hours). N = 6 independent wells are shown for each the vehicle and rotenone conditions. Bars represent mean, error bars represent standard error of the mean. P values were determined using Student’s t-test. **(G)** ROS levels (CellROX^TM^ orange staining) comparing 2D monoculture neurons versus 3D monoculture neurons. N=3 wells per sample. Bars represent means of the median fluorescence level and error bars represent standard error of the mean. P values were determined using Student’s t-test. **(H,I)** ROS levels CellROX^TM^ orange staining). comparing 3D monoculture neurons with neuron-astrocyte co-cultures of variable ratios. Bars represent normalized mean ROS levels compared to 3D monoculture neurons. N= 3 independent wells. Error bars represent standard error of the mean . ANOVA followed by Tukey’s multiple comparison test was conducted to determine significance. **(J, K)** Peroxidated lipid levels of *GSK3*Β KD cells as compared to NTC. Cells were stained with Liperfluo and median fluorescence levels were measured. Levels were then normalized to the NTC. J, example trace. K, quantification of 5 replicates (5 cell culture wells), bars represent mean, error bars represent standard error of the mean . P values were determined using Student’s t-test. **(L)** Reactive oxygen species (ROS) levels of 3D neurons treated with tetrodotoxin (TTX, 1.5 µM, duration of 1 week). Neurons are stained with CellROX orange and fluorescence levels were read out using flow cytometry. median fluorescence was normalized to the fluorescence level of the vehicle condition to account for different levels of background fluorescence across experiments. Dots represent n = 4 individual wells. Error bars represent standard error of the mean . P values were determined using Student’s t-test. **(M)** Peroxidated lipid levels (Liperfluo stain) in 3D monocultured neurons treated with vehicle, tetrodotoxin (1.5 µM, duration of 1 week), and ferrostatin (10 µM, duration of 1 week). Bars represent mean, error bars represent standard error of the mean. Dots represent cells from n = 3 independent wells. ANOVA followed by Tukey’s multiple comparison test was conducted to determine significance. **(N)** Viability of 3D monocultured neurons after 14 days in culture with and without tetrodotoxin (TTX, 1.5 µM, duration of 1 week) and ferrostatin (10 µM, duration of 1 week) treatment. Cells were stained with trypan blue, and the percentage of trypan blue negative cells were obtained. Bars represent mean, error bars represent. Error bars represent standard error of the mean . N=7 wells for vehicle, 8 wells for ferrostatin and 6 wells for TTX. ANOVA followed by Tukey’s multiple comparison test was conducted to determine significance.

Upon induction of oxidative stress, NRF2 translocates to the nucleus and activates transcription of genes protective against oxidative stress^36,40,41^. To characterize how NRF2 activity may be modified by *GSK3B* knockdown, we sectioned iAssembloids into 20-micron sections and determined the levels of nuclear NRF2. We found that knockdown of *GSK3B* resulted in higher levels of NRF2 in the nucleus (Fig. 5C), consistent with the induction of its target genes (Fig. 4C). We then asked whether the knockdown of *GSK3B* itself increased oxidative stress, resulting in the translocation of NRF2 to the nucleus. To test this, we assessed reactive oxygen species (ROS) levels by using the dye, CellROX^TM^ Orange. Knockdown of *GSK3B* did not influence the level of ROS in neurons in 2D or 3D culture (Fig. 5D, Fig. S6A). This suggests that knockdown of *GSK3B* does not drive oxidative stress and that *GSK3B* knockdown activates NRF2 is downstream of oxidative stress.

GSK3 activity has been reported to be activated in response to ROS to inhibit NRF2 by preventing its translocation to the nucleus in a model for acute kidney injury as well as in cortical neurons^42–45^. We hypothesized that GSK3 activity prevented neurons from mounting an appropriate response to oxidative stress using a similar mechanism. To test this hypothesis, we first assessed differences in GSK3 expression and activity in 2D neurons versus 3D neurons and iAssembloids. We found that although *GSK3B* expression did not differ between culture systems, GSK3 activity (denoted by the loss of the inhibitory phosphorylation site at serine 9), was 3-4 times higher in 3D culture versus 2D neurons (Fig. 5E). We asked if oxidative stress could induce this change in GSK3 activity, and indeed, by adding rotenone to 2D neurons, which increases ROS levels (Fig. S6B), we found that the activity of GSK3 increased (Fig. 5F). We then assessed ROS levels in 2D as well as 3D neurons and found that ROS levels in 3D neurons were consistently higher than in 2D (Fig. 5G, Fig. S6C). This suggests that GSK3 activity prevents the NRF2-mediated protective response to oxidative stress, and that knockdown of GSK3 confers a beneficial effect for cell survival by increasing the NRF2 response. We then asked what the role of glial cells such as astrocytes play in this response.

We found that by supplementing more glia, we were able to decrease the level of ROS in the neurons (Fig. 5H-I). We then asked whether knockdown of *GSK3B* could ameliorate downstream effects of increased ROS, such as lipid peroxidation^7^. Indeed, we found that knocking down *GSK3B* led to a decrease in the level of peroxidated lipids in neurons (Fig. 5J-K, Fig. S6D). Peroxidated lipids can drive neuronal death via ferroptosis^7,46^. Therefore, we hypothesized that the neuronal death occurring in 3D culture was due to this accumulation of peroxidated lipids driving ferroptosis.

Since neuronal activity is much higher in iAssembloids compared to 2D-cultured neurons, and level of activity correlates with proportion of glial cells (Fig. 2E), we hypothesized that neuronal activity could be driving higher levels of ROS and thus ferroptosis induced cell death. We found that ROS levels of neurons in iAssembloids which contains glia are lower when compared to those of 3D neuronal monocultures (Fig. S6E). Interestingly, neuronal activity is much higher in 3D-cultured neurons compared to iAssembloids, which contain glial cells, suggesting that glial cells may prevent neuronal hyperexcitability, such as through glutamate-buffering or through ROS clearing activity of astrocytes^47–49^. Previous studies have shown that high levels of neuronal activity lead to a generation of ROS, which can drive excitotoxicity^50,51^. To test whether neuronal activity drives the increase in ROS in our culture system, we added tetrodotoxin (TTX), a drug that blocks voltage-gated sodium channels and thus neuronal firing. We found that TTX treatment decreased ROS levels (Fig. 5L) and levels of peroxidated lipids, to similar levels as the ferroptosis blocker ferrostatin. (Fig. 5M). We also found that blocking neuronal activity and ferroptosis with ferrostatin improved neuronal survival in 3D neurons (Fig. 5N).

Based on the data above, we hypothesize that in 2D-monocultured neurons, there is a low level of neuronal activity and ROS levels remain low, thus neurons are not at risk for death via ferroptosis (Fig. 6A). However, in 3D monocultured neurons, higher levels of neuronal activity trigger a build-up in ROS, and both processes increase GSK3 activity. This increase in GSK3 activity blocks NRF2 from protecting neurons from oxidative stress, ultimately leading to neuronal death via ferroptosis (Fig. 6B). In GSK3 knockdown neurons, the ability for GSK3 to block NRF2 from protecting neurons from neuronal death is abrogated, enabling the protective NRF2-mediated oxidative stress response that improves neuronal survival (Fig. 6C). The presence of glial cells in iAssembloids partially reduces neuronal activity and ROS in neurons, thereby reducing, but not abrogating GSK3 activity and its suppression of protective NRF2 activity (Fig. 6D).

**Figure 6.**
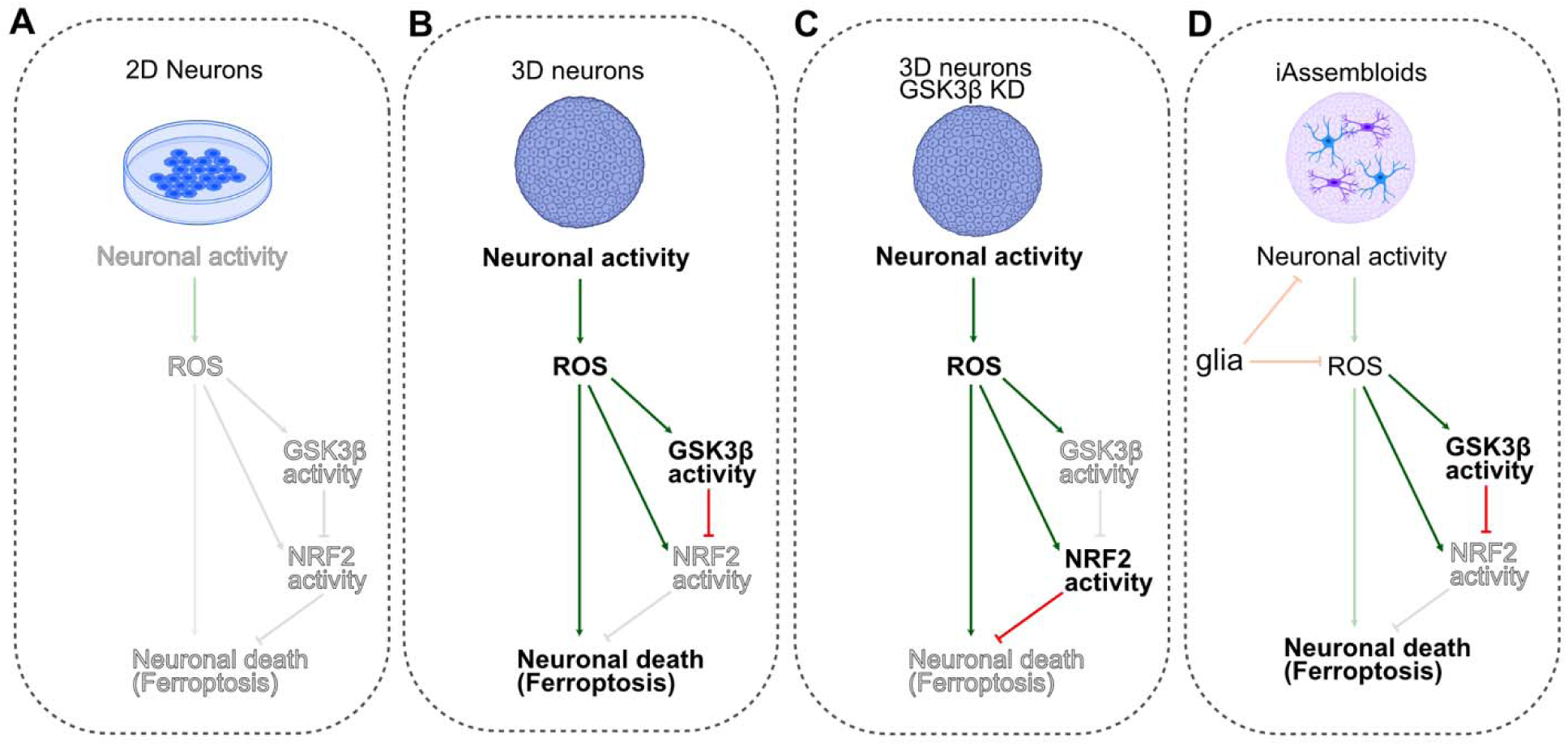
Proposed model of GSK3 function in oxidative stress and neuronal death **(A)** In 2D monocultured neurons, there is not sufficient neuronal activity to drive oxidative stress. Therefore, GSK3 activity is low and neuronal death is not triggered. **(B)** In 3D monocultured neurons, neuronal activity is high. This triggers an increase in ROS and oxidative stress and leads to an increase in GSK3 activity. Increase of GSK3 activity blocks NRF2 from translocating to nucleus to mount a protective oxidative stress response. In the absence of the protective response, neurons undergo ferroptosis. **(C)** In 3D monocultured neurons with *GSK3B* knockdown, neuronal activity and oxidative stress is triggered, but in the absence of GSK3 , NRF2 can translocate to the nucleus, preventing ferroptosis. **(D)** In iAssembloids, the addition of glial cells reduces neuronal activity and ROS, possibly by regulation of glutamate levels via glutamate reuptake, which could lessen GSK3 activity and protect cells against ferroptosis.

### CRISPRi screens in iAssembloids uncover cell non-autonomous effects of astrocytes expressing different Apolipoprotein E (APOE) variants on neuronal health

We next applied the iAssembloid system to study the effects of glial cells on neurons in a neurodegenerative disease-relevant context. We were particularly interested in the genetic risk factor for Alzheimer’s Disease, APOE- 4. APOE- 4 is a variant of the APOE gene and carriers of the APOE- 4 allele are much more likely to develop Alzheimer’s Disease^52,53^. The APOE- 3 allele is considered “neutral.” APOE is mainly expressed by glial cells such as astrocytes, although stress can induce its expression in neurons^54,55^. However, how expression of these APOE variants in glial cells may affect neuronal survival is currently unknown.

To assess this, we designed a CRISPRi-based screen in which we transduced ∼16,000 sgRNAs targeting ∼2,000 genes in the “druggable genome” (H1 library)^27^ into hiPSCs expressing dox-inducible NGN2 and CRISPRi machinery. These neurons are homozygous for the APOE- 3 allele. We then differentiated these cells into neurons and co-cultured them with either APOE- 3 homozygous or APOE- 4 homozygous astrocytes for 14 days. We then sequenced the sgRNAs from the surviving neurons to determine the sgRNA frequencies in the starting versus surviving population in both conditions. By comparing the hits from the APOE- 4 and the APOE- 3 screen, we can uncover pathways in neurons that differentially control their survival in the presence of APOE- 4 and the APOE- 3 astrocytes (Fig. 7A).

**Figure 7.**
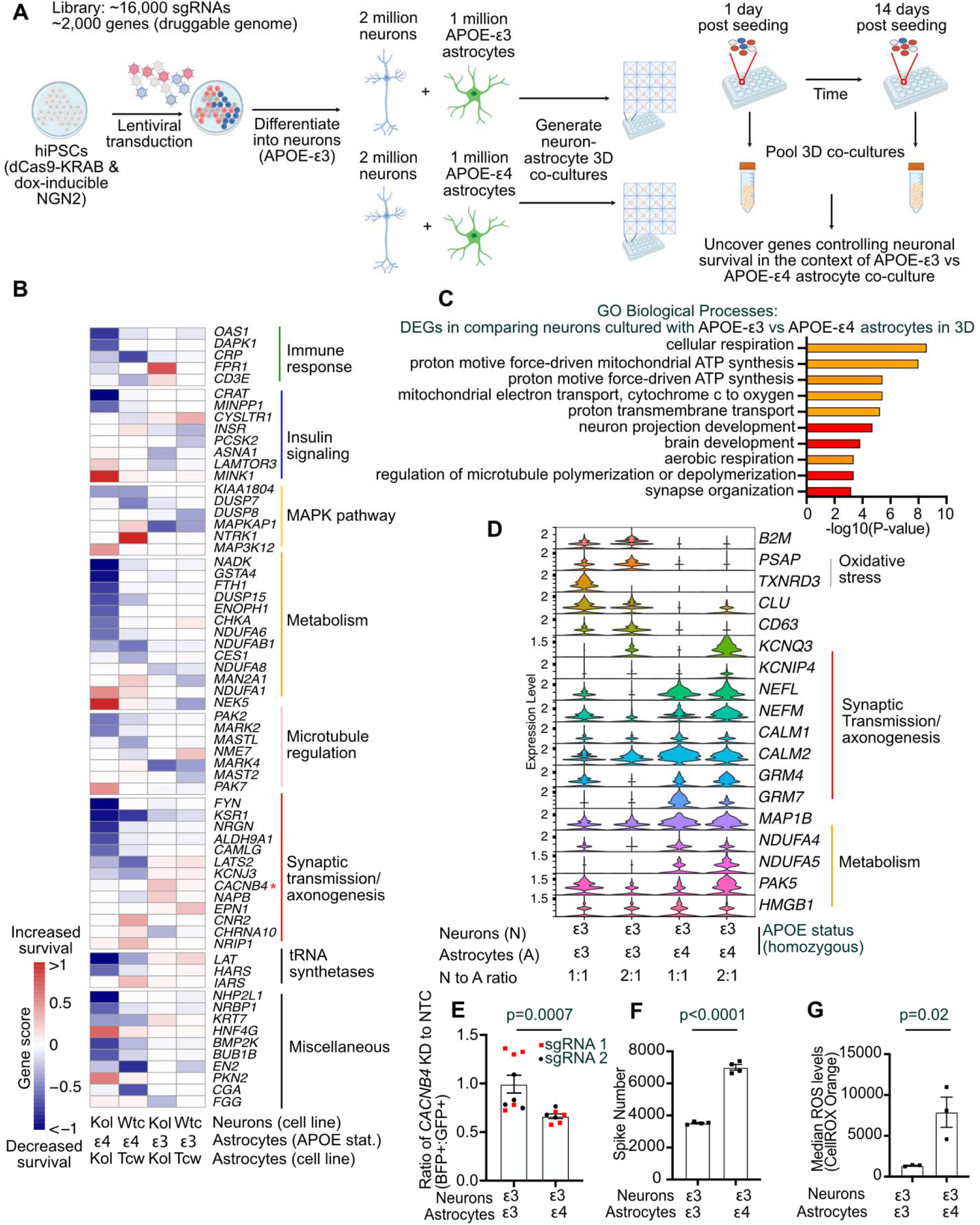
CRISPRi-based screen for neuronal survival in APOE-ε3 versus APOE-ε4 astrocyte 3D co-cultures **(A)** Schematic of experimental design. Screens for neuronal survival were conducted in APOE- ε3 neurons expressing dCas9-KRAB and dox-inducible CRISPRi that were 3D co-cultured with APOE-ε3 vs APOE-ε4 astrocytes. **(B)** Hit genes that were consistent between the two screens (WTC11 and KOLF2.1) and had the strongest difference in gene score between APOE-ε3 and APOE-ε4 astrocyte co-cultures were selected and annotated. Heatmap represents gene scores. **(C)** Top 10 enriched GO biological processes differentially expressed genes in neurons 3D-co- cultured with APOE-ε3 versus APOE4-ε4 astrocytes, based on snRNA-seq. Orange bars represent terms related to metabolic processes and red bars represent terms related to neuronal function. P-values are calculated by EASE Score, a Modified Fisher Exact P-value. **(D)** Violin plot of expression of genes selected from related GO terms across different co-culture conditions from the snRNA-seq dataset represented in C. **(E)** Validation of screening results of *CACNB4* KD. Neurons expressing non-targeting control guides and GFP were seeded at 1 to 1 ratio with neurons expressing sgRNAs targeting *CACNB4* and BFP. Neurons were then cocultured with APOE-ε3 astrocytes (n=5 wells for sgRNA 1, n=4 wells for sgRNA 2) vs APOE-ε4 astrocytes (n=4 wells for sgRNA 1, n=4 wells for sgRNA 2) and the ratio of BFP+ to GFP+ cells were assessed after 2 weeks. P-value from Student’s t-test. Error bars represent standard error of the mean . **(F)** Multi-electrode array analysis of neurons co-cultured with APOE-ε3 vs APOE4-ε4 astrocytes. Spike numbers are represented. n=4 wells for each condition. P-value from Student’s t-test. Error bars represent standard error of the mean . **(G)** Median CellRox Orange fluorescence levels for neurons co-cultured with APOE-ε3 vs APOE4-ε4 astrocytes. n=3 wells for each condition. P-value from Student’s t-test. sError bars represent standard error of the mean .

We conducted this screen in two different cell line background pairs: one where the neurons and astrocytes were not isogenic, with astrocytes generated from the TCW 1E33 and 1E44 background^56^, and neurons from WTC11 background^6^ (Fig. S7A, Table S5), and another where the neurons and astrocytes were isogenic with cells from the iNDI project (KOLF2.1)^57^ (Fig. S7B, Table S5). We found that hits obtained from both screens were similar, with pathways that were related to insulin signaling, MAPK pathway, metabolism, immune response, microtubule regulation, and synaptic transmission most consistently different between the APOE- 3 and APOE- 4 conditions (Fig. 7B). However, we focused our investigations onto the KOLF2.1 line due to recent information suggesting some karyotypic abnormalities in the TCW line (personal communication, Julia TCW). To further investigate differences between the culture systems, we conducted single-nucleus RNA sequencing on nuclei isolated from the KOLF2.1 3D neuron-astrocyte co-cultures in with neurons and astrocytes at 1 to 1 or 2 to 1 ratio (Fig. S7C). We separated the neurons and astrocytes based on cell type-specific markers and focused mainly on neuronal gene expression (Fig. S7D-E). In general, many of the differentially expressed genes fell into categories relating to oxidative phosphorylation/metabolism or neuronal functions such as synaptic organization (Fig 7C, Table S6). We found that genes such as *B2M, PSAP, TXNBD3, and CLU* were more highly expressed in neurons co-cultured with APOE- 3 astrocytes while other genes related to synaptic transmission and axon development (e.g. *GRM4, GRM7, NEFL, NEFM, KCNQ3)* were more highly expressed in neurons co-cultured with APOE- 4 astrocytes (Figure 7D). To further investigate the connection between synaptic functions in our APOE- 3 vs APOE- 4 co-culture systems, we focused on one of our genes of interest from our screen, *CACNB4 (*Calcium Voltage-Gated Channel Auxiliary Subunit Beta 4).

*CACNB4* encodes the beta subunit of voltage-gated calcium channels located in the synapse. This subunit is important for regulating the properties of calcium currents by modulating the gating properties of the channel. Loss of function in this subunit has been associated with epilepsy^58^. In our screen, we found that knockdown of CACNB4 leads to a stronger deficit in neuronal survival in co-culture with APOE- 4 astrocytes as compared to co-culture with APOE 3 astrocyte (Fig. 7B). We validated this phenotype by co-culturing neurons expressing a non-targeting GFP+ sgRNA and a *CACNB4* BFP+ targeting guide with either APOE- 3 or APOE- 4 expressing astrocytes. We then assessed the ratio of BFP+ to GFP+ cells after 2 weeks in culture. We found that indeed, knockdown of *CACNB4* had a stronger effect on neuronal survival in co-culture with APOE- 4 expressing astrocytes as opposed to APOE- 3 expressing astrocytes (Fig. 7E). Therefore, we hypothesized that there may be differences in neuronal activity in these two culture systems, leading to the differential accumulation of ROS. We performed multi-electrode array analysis to assess whether neuronal activity differed between the APOE- 3 and APOE- 4 conditions. Indeed, neurons co-cultured with APOE- 4 astrocytes had higher spike rates than neurons co-cultured with APOE- 3 astrocytes (Fig. 7F). This correlated with ROS levels detected, as neurons co-cultured with APOE- 4 astrocytes had higher levels of fluorescence of the CellROX orange dye compared to neurons co-cultured with APOE- 3 astrocytes (Fig 7G). Thus, we hypothesize that APOE- 4 expressing astrocytes may differentially affect neuronal activity as compared to APOE- 3 expressing astrocytes, leading to increased sensitivity to neuronal death in APOE- 4 co-culture systems.

In our dataset, we had found lower expression of *CACNB4* in neurons co-cultured with APOE-4 astrocytes (Table S6). We hypothesized that the decrease in *CACNB4* expression may be due to the drop out of a specific subset of neurons in the APOE- 4 environment. We examined whether APOE- 3 vs 4 genotypes would affect expression levels of *CACNB4* in human AD patient brains, based on a previously published single-nuclei RNA sequencing dataset by Haney et al^26^. To do this, we subset excitatory (*RBFOX3+*, *SLC17A7+*) neurons in the human AD dataset by Haney et al^26^. We then performed unbiased clustering of this population of cells, and we found that clusters 1 and 7 consisted of cells with lower *CACNB4* expression (Fig S7F).

Through differential expression analysis, we identified markers for these clusters. We found that cells within these clusters also expressed other markers such as *CUX2* for cluster 1, and *RELN* for cluster 7 (Fig S7E, Table S6). *CUX2* (Cux2 cut-like homeobox 2) is a transcription factor that is highly expressed in upper layers of the brain, particularly in the pyramidal neurons of layer II-IV. Interestingly, *RELN* has been proposed to be a marker of selective vulnerable neurons ^59,60^. In these clusters, we find that clusters 1 contain a higher proportion of neurons from patients with no disease as opposed to those diagnosed with Alzheimer’s disease (Fig. S7F-G). Interestingly, cluster 7 had similar proportions of neurons from APOE- 3 patients diagnosed with and without disease. For both these clusters, we also found that these clusters contain more APOE- 3 neurons than APOE- 4 AD patient neurons (Fig. S7G). These findings are compatible with the hypothesis that low levels of *CACNB4* expression contribute to the vulnerability of specific neuronal subpopulations in AD, in particular in an APOE- 4 background.

## Discussion

In this study, we use a scalable 3D co-culture model, which we call iAssembloids, in combination with CRISPRi-based functional genomics screens to uncover how genes regulate neuron and glia function and interactions in a scalable manner. iAssembloids are fast to generate and homogeneous, which is ideal for CRISPR-based screens and other systems requiring high throughput screening (e.g., compound screening). Relative to monoculture, neurons, and glia in iAssembloids adopt more mature states. This enabled us to uncover a pathway controlling the survival of highly active neurons, which would have been difficult to do in 2D monoculture neurons. iAssembloids allow precise control of glial cell genetic backgrounds, allowing causal assessment of those genetic backgrounds on neuronal survival. We demonstrate this application to examine the effect of glia-expressed APOE-ε4, a variant of the gene *APOE* known to increase risk of late-onset Alzheimer’s Disease^52,53^, on neuronal survival.

### GSK3β as a player in oxidative stress in response to neuronal activity

Many studies have reported a link between seizure activity and dementia in advanced stages^61–63^. However, developing unprovoked seizures later in life have also been associated with early-stage dementia and have been posited as one of the first signs of neurodegenerative disease^64^. Our model shows that neurons in iAssembloids are more electrophysiologically active than in monoculture, thus allowing us to assess the pathways that are affected by neuronal activity in neuronal survival.

Our CRISPRi-based screens uncovered GSK3β as a factor controlling the response to neuronal activity and oxidative stress. In Alzheimer’s Disease, GSK3β plays a major role in phosphorylating Tau, which is thought to contribute to Tau aggregation^34,65^. It is also commonly found to be hyperactivated in brains of Alzheimer’s Disease patients^30,66^. Here, we unravel a secondary role for GSK3β that may contribute to its role in neurodegeneration. In this case, neuronal activity, and an increase in reactive oxygen species (ROS) can drive GSK3β activity.

GSK3β blocks NRF2 from mounting a protective response to oxidative stress. We also find that the addition of glial cells can be protective against the generation of reactive oxygen species, which is consistent with previously described roles of glia in response to neuronal activity and oxidative stress^47,48^. Therefore, we hypothesize that in disease, the combined effects of aberrant neuronal activity driving increases in ROS and GSK3β activity as well loss of glial cell homeostasis in protecting neurons against ROS can drive neurodegeneration.

Our findings suggest that there may be some beneficial effects of GSK3β inhibition in neurons that enables the activation of a protective response against oxidative stress in excitotoxic conditions. GSK3β is a kinase that has many downstream targets, making it difficult to target as a therapeutic target^67^. However, more specific therapeutic strategies to boost the oxidative stress response could recapitulate some beneficial aspects of GSK3β inhibition, without off-target effects.

Beyond GSK3β, we also observe other genetic perturbations, such as knockdown of *CDK8* and *MAPK1*, that can yield similar transcriptional responses based on our CROP-seq data set. Future studies will examine whether these genes act through a similar pathway as GSK3β, or through an alternative pathway.

### APOE-ε3 vs APOE-ε4 screen suggest differences in synaptic transmission in the co-culture system

In our screen comparing neuronal co-cultures with APOE-ε3 versus APOE-ε4 astrocytes, we found many differential targets relating to insulin signaling, the MAPK pathway, and most strikingly, synaptic transmission. We hypothesize that APOE- 4-expressing astrocytes may be linked to neuronal hyperexcitability either by gain of toxic or loss of protective functions. Recent studies have suggested that hyperactivity can precede neuronal degeneration in the human Alzheimer’s disease brain^68^. Conversely, aberrant neuronal activity can also have a detrimental effect on surrounding glial cells, such as microglia. However, it was shown that co-culture with APOE- 4 microglia decreases the level of neuronal network activity in a spheroid system relative to co-culture with APOE- 3 microglia, due to microglial dysfunction^69^. Interestingly, we find that APOE- 4 astrocytes have an opposing effect in that APOE- 4 astrocyte co-culture leads to an overall higher level of neuronal activity in our spheroid system relative to co-culture with APOE- 4 astrocytes. Although the net result on neuronal activity is opposing, both studies highlight that APOE- 4 expression in glial cells can contribute to alterations in neuronal activity. In *vivo*, young APOE- 4 KI mice have also been found to have hippocampal region-specific network hyperexcitability, supporting our finding that an APOE- 4 cellular environment may lead to hyperactivity in neurons ^70^. Based on our study, APOE- 4 astrocytes may exacerbate neuronal loss by promoting neuronal hyperactivity and driving ROS accumulation in neurons.

Future studies will focus on understanding the mechanism by which in APOE-ε4 astrocytes may drive neuronal hyperactivity.

### Neuronal activity as a link between *GSK3B and APOE*

Our system highlights the possibility of neuronal hyperactivity as a link between neuronal death driven by GSK3B activity as well as APOE-ε4 genotype. APOE-ε4 astrocytes increase neuronal activity, driving ROS and an increased risk of ferroptosis, and GSK3B activation prevents the effective mounting of a protective oxidative stress response. Interestingly, patient studies have shown that GSK3B activity in addition with the APOE-ε4 genotype have been associated with cognitive decline in patients with type 2 diabetes. These patients then have a higher incidence of developing Alzheimer’s Disease^71^. Future research can further elucidate the possible pathways that may connect the two.

### Limitations of this study

Our model does not recapitulate all aspects of an *in vivo* brain environment. It is a simplified model that does not mimic the vast amount of cellular heterogeneity found inside a human brain. For instance, iAssembloids do not contain vasculature or other glial cell types, such as oligodendrocytes. It also does not seek to mimic potential region-specific effects that differ from brain region to brain region. Due to the nature of the NGN2-induced neurons we use, we also cannot accurately model aspects of brain development or tissue architecture. Other systems use methods to generate brain organoids that more closely aligns with development, but these methods can be more labor and time intensive^11,12,72^. Our platform seeks to address questions on a cellular level, enabling mixing and matching neuronal and glia subtypes using a reductionist approach, thus pinpointing the specific contributions to a phenotype by a specific glial subtype. It also enables the examination of non-cell autonomous contributions of specific genetic variants on neuronal function, such as APOE-ε4. The model can be expanded upon by integrating vasculature as previously shown by others^73,74^ and can be applied to study the glial contributions of other genetic risk factors.

Neuronal survival is highly relevant phenotype for neurodegenerative diseases. However, it is a late phenotype that is the result of many preceding functional changes a neuron may experience. Future studies will investigate the functional effects of genetic perturbations such as GSK3β on more complex phenotypes such as neuronal activity, states of glial cells, and intercellular connections. Beyond these studies, this model could provide an additional platform to assess the effectiveness of small molecules on neuronal phenotypes relating to neuronal activity. In the future, this model could also provide a unique platform to examine cell-cell interactions in the context of other neurological diseases where neuron-glia interactions play an important role, such as other neuropsychiatric and neurodegenerative disorders.

## Acknowledgements

We would like to thank Noam Teyssier for helpful discussions relating to single-cell analysis. We thank Dr. Kun Leng, Sydney Sattler, for their advice regarding model generation. We thank Greg Mohl, Drs. Lisa Boxer, Amanda McQuade, William Renthal and Marty Yang for manuscript suggestions. We thank Stephanie Huard, Anjani Atilli, Dr. Marty Yang, Lydia Lee, Merissa Chen, and Jason Hong for technical support. We thank David Shin and Jenelle Wallace for their advice regarding multi-electrode array analysis, Rene Sit, Michelle Tan, and Norma Neff at CZ Biohub for help with sequencing, and Yoshi Sei, Neal Bennett, and the Nakamura Lab for helpful discussions. For reagents, we would like to thank the contributions of Dr. Alison Goate and Dr. Julia TCW and Bill Skarnes and the iNDI project. We thank past and present members of the Kampmann Lab for their support and guidance in this project.

The authors would like to thank Sarah Elmes and team at the UCSF Laboratory for Cell Analysis, Eric Chow, and team at UCSF Center for Advanced Technology, and finally, Kari Harrington and DeLaine Larsen at the Nikon Imaging Core and Weill Imaging Cores for their advice and assistance.

Images of neurons, astrocytes, microglia and 3D cultures were generated using Biorender and modified for figures.

M. Kampmann was supported by NIH/NIA U01 AG072464, NIH/NINDS U54 NS123746, The Tau Consortium / Rainwater Charitable Foundation, and a Ben Barres Early Career Acceleration Award from the Chan Zuckerberg Neurodegeneration Challenge Network. E.L. was funded by the Department of Defense National Defense Science and Engineering Graduate Fellowship and the UCSF Fletcher Jones Fellowship. C.B. is supported by the NSF Graduate Fellowship. S.C.B is supported by the Alzheimer’s Association Research Fellowship Program (23AARF-1027616). O.M.T. is funded by the National Science Foundation Graduate Research Fellowship under Grant No. 2034836. I.V.L.R. is funded by California Institute for Regenerative Medicine (CIRM) grant EDUC412812 and NIH grant T32NS115706. A.J.S was supported by NIH grant F32AG063487 and K99AG080116-01. PIP-seq sequencing was performed at the UCSF CAT, supported by UCSF PBBR, RRP IMIA, and NIH 1S10OD028511-01 grants.

## Author contributions

E.L. and M. Kampmann contributed to the study’s overall conception, design, and interpretation and wrote the manuscript and created the figures with input from the other authors. E.L. optimized the iAssembloid model with helpful discussions from M. Koontz and E.M.U. E.L. characterized the iAssembloid model, performed and analyzed the CRISPRi-based screens (including survival screens and CROP-seq), and performed mechanistic follow-up on relevant targets with guidance from M. Kampmann. N.D. and M. Kampmann. selected genes for and cloned the neurodegeneration library. M. Koontz supplied serum-free astrocytes for CRISPRi-based screens. D.M. provided support in analyzing and single cell sequencing of APOE3 and APOE4 co-culture. C.B. and E.L. generated data from multi-well multi-electrode arrays in iAssembloids 3D co-cultures and S.C.B. generated corresponding data in 2D monoculture neurons. A.J.S. performed western blots in rotenone induced 2D monocultured neurons.

I.V.L.R. and E.L. generated APOE3 and APOE4 KOLF2.1 astrocytes used in this study. O.M.T. generated 6TF hiPSC cell line expressing mScarlet. N.P. cloned sgRNAs utilized for APOE3 and APOE4 validation experiment.

## Declarations of interest

M.K. is a co-scientific founder of Montara Therapeutics and serves on the Scientific Advisory Boards of Engine Biosciences, Casma Therapeutics, Alector, Montara Therapeutics and Neurocrine, and is an advisor to Modulo Bio and Recursion Therapeutics. M.K. is an inventor on US Patent 11,254,933 related to CRISPRi and CRISPRa screening, and on a US Patent application on in vivo screening methods.

## Methods

### hiPSC Maintenance and Culture

We cultured hiPSCs in the WTC11^75^ and KOLF2.1^57^ backgrounds on plates coated with Matrigel (Corning, Cat. No. 356231) at a concentration of 1:100 in Knockout DMEM (Gibco/Thermo Fisher Scientific, Cat. No. 10829-018). For plating, hiPSCs were seeded with 10 µM Rock Inhibitor (Y-27632, Biotechne/Tocris Cat. No. 1254) diluted in Gibco™ StemFlex™ Medium (Gibco/Thermo Fisher Scientific, Cat. No. A3349401). Media (no RI) was exchanged when colonies reached ∼20 cells in size. hiPSCs were maintained with media changes every day.

When hiPSCs reached ∼80% confluence, hiPSCs were coated with either Gibco™ Versene Solution (0.48 mM) (Gibco/Thermo Fisher Scientific, Cat. No. 15040066) for routine passaging or StemPro™ Accutase™ Cell Dissociation Reagent (Gibco/Thermo Fisher Scientific, Cat. No. A1110501) for differentiation and incubated at 37 °C for 5-7 minutes. After dissociation, cells were collected and rinsed with 1X DPBS (MilliporeSigma, Cat. No. D8537). Cells were centrifuged at 250g for 5 minutes and replated.

### hiPSC derived Neurons

hiPSC-derived neurons were differentiated using methods previously described^6,7^. Briefly, we dissociated hiPSCs containing doxycycline-inducible NGN2 in the AAVS1 locus and pC13N- dCas9-BFP-KRAB in the CLYBL locus and re-plated cells on Matrigel coated plates with Pre-differentiation medium containing Knockout DMEM/F12 (Gibco/Thermo Fisher Scientific, Cat. No. 12660-012), 1X GlutaMAX Supplement (Gibco/Thermo Fisher Scientific, Cat. No. 35050- 061), 1X MEM Non-Essential Amino Acids (Gibco/Thermo Fisher Scientific, Cat. No. 11140- 050), 1X N2 Supplement (Gibco/Thermo Fisher Scientific, Cat. No. 17502-048), 10uM ROCK inhibitor (Y-27632, Biotechne/Tocris, Cat. No. 1254), 1 µg/mL mouse laminin (Thermo Fisher Scientific, 23017-015), 10 ng/mL BDNF (PeproTech, Cat. No. 450-02B) and 10 ng/mL NT3 (PeproTech, Cat. No. 450-03B) with 2µg/mL doxycycline (Takara Bio, Cat. No. 631311) to induce expression of NGN2. Half of the media was exchanged over the next 2 days until hiPSC-derived neurons were ready for iAssembloid generation (Day 0). For monocultures, pre-differentiated neurons were plated onto PDL-coated plates (Corning, Cat. No. 356469) and maintained in differentiation media: Base media: 50% DMEM/F12 (Gibco/Thermo Fisher Scientific, Cat. No. 11320-033) and 50% Neurobasal-A (Gibco/Thermo Fisher Scientific, Cat.

No. 10888-022) with supplements: 1X MEM Non-Essential Amino Acids, 0.5X GlutaMAX Supplement, 0.5X N2 Supplement, 0.5X B27 Supplement (Gibco/Thermo Fisher Scientific, Cat. No. 17504-044), 10 ng/mL NT-3, 10 ng/mL BDNF and 1 μg/mL Mouse Laminin.

### hiPSC derived Astrocytes

#### Serum Free Protocol

Astrocytes were generated as described previously^13,14,16^. WTC11 iPSCs were cultured and expanded in six-well plates using Essential 8 media (Gibco/Thermo Fisher Scientific, Cat. No. A1517001). Once iPSCs were confluent, cells were enzymatically dissociated into a single cell suspension using Accutase (Gibco/Thermo Fisher Scientific, Cat. No. A1110501). The resulting single cell suspension was transferred into T-75 flasks in ∼10 ml of neural induction media (50% DMEM/F12 (Gibco/Thermo Fisher Scientific, Cat. No. 11320-033) and 50% Neurobasal-A (Gibco/Thermo Fisher Scientific, Cat. No. 10888-022) with supplements: 1X MEM Non-Essential Amino Acids (Gibco/Thermo Fisher Scientific, 11140050), 1X GlutaMAX Supplement (Gibco/Thermo Fisher Scientific, Cat. No. 35050-061), 1X N2 Supplement (Gibco/Thermo Fisher Scientific, Cat. No. 17502-048), 0.5X B27 Supplement (Gibco/Thermo Fisher Scientific, Cat. No. 17504-044), Anti-Anti (Gibco/Thermo Fisher Scientific, Cat. No.15240062) to allow iPSCs to aggregate into small spheroids in suspension. To facilitate neural induction, dual SMAD inhibitors DMH1 (Tocris, Cat. No. 73634) and SB-431542 (StemCell Technologies, Cat. No. 72234) are added to the media. Once rosette formation is observed, spheroids are transferred to culture dishes coated with Matrigel (Corning, Cat. No. CB40230C) and allowed to attach and distributed across culture dish surfaces. Once sufficiently distributed, neuroepithelial rosettes are mechanically removed and isolated. Neuroepithelial spheroids are maintained is suspension in neural media (DMEM-F12, N2/B27 supplements, Heparin (Stemcell Tech, Cat. No. 07980), Anti-Anti) plus mitogenic growth factors EGF (Peprotech, Cat. No. AF-100-15) and FGFβ (Peprotech, Cat. No. 100-18B). Spheroid cultures are maintained in suspension and triturated/dissociated regularly to maintain size and structural homogeneity of neuroepithelial spheroids. After ∼180 days of culture maintenance, neuroepithelial progenitor cells have differentiated into unipotent populations of glial progenitors.

#### Serum Based Protocol

Astrocytes were differentiated as described previously^56^. hiPSCs were cultured on Matrigel and in StemFlex media supplemented with 10 μM RI. hiPSCs were then differentiated to neural precursor cells by dual SMAD inhibition using 0.1 mM LDN193189 (Biotechne/Tocris Cat. No. 6053) and 10 mM SB431542 (Biotechne/Tocris, Cat. No. 1614) in N2/B27 media inhibition in embryoid bodies (EB) media (DMEM/F12 + 1x N2 + 1x B27-VA). EBs were cultured in AggreWell™800 (StemCell Technologies, Cat. No. 34815) plates rinsed with Anti-Adherence solution (StemCell Technologies, Cat. No. 07010) for 7 days and then plated onto a Matrigel coated plate. Neural rosette formation occurs at day 14 where rosettes were selected for with STEMdiff™ Neural Rosette Selection Reagent (StemCell Technologies, Cat. No. 05832). NPCs were expanded in NPC media which contained DMEM/F12 + 1x N2 + 1x B27-VA supplemented with FGF2 (Biotechne, Cat. No. 233-FB-010). Using Magnetic Activated Cell Sorting (MACS).

NPCs were enriched by sorting against neural crest cells (Miltenyi Biotech, Cat. No. 130-097- 127) and for CD133+ cells (Miltenyi Biotech, Cat. No. 130-097-049) using LD (Miltenyi Biotech, Cat. No. 130-042-901) and LS (Miltenyi Biotech, Cat. No. 130-042-401) columns respectively. Magnet was 3D printed using previously published protocol^76^. by NPCs were maintained in NPC media (were validated with immunostaining for SOX2 (Rabbit anti-SOX2, Abcam, Cat. No. ab97959) and Nestin (rabbit anti-Nestin, Abcam, Cat. No. ab92391). Dissociated NPCs were transferred to astrocyte media (ScienCell, Cat. No. 1801) and cultured for 30 days on Matrigel until they stained positively for S100B (mouse anti-S100B, MilliporeSigma, Cat. No. S2532) and NFIA (MilliporeSigma, Cat. No. HPA008884).

### hiPSC derived Microglia

Microglia were differentiated as previously described. Briefly, hiPSCs expressing six inducible transcription factors (MAFB, CEBPα, IRF, PU1, CEBPβ, IRF5) were grown in Essential 8™ Basal Medium (Gibco/Thermo Fisher Scientific, Cat. No. A15169-01), 10 μM ROCK inhibitor, and 2 μg/ml Doxycycline on Matrigel and 10 cm Poly-D-Lysine coated plates at a density of 1.5 million cells per dish. After 2 days, cells were grown in a differentiation media consisting of Advanced DMEM/F12 (Gibco/Thermo Fisher Scientific, Cat. No. 35050-061), 1X GlutaMAX, 2ug/ml doxycycline, 100 ng/mL Human IL34 (Peprotech; Cat. No. 200-34) and 10 ng/mL Human GM-CSF (Peprotech; Cat. No. 300-03). Two days later, media was exchanged for iTF-Microglia media consisting of Advanced DMEM/F12, 1X GlutaMAX, 2 μg/mL doxycycline, 100 ng/mL Human IL-34, 10 ng/mL Human GM-CSF, 50 ng/mL Human M-CSF (Peprotech; Cat. No. 300- 25) and 50 ng/mL Human TGFB1 (Peprotech; Cat. No. 100-21C). On day 8, cells were dissociated with TypLE express (Gibco/Thermo Fisher Scientific, Cat. No. 12605-028) and seeded into iAssembloids.

## Generation of iAssembloids

AggreWell™800 plates were first rinsed with Anti-Adherence rinsing solution and then washed with DMEM/F12. Neurons (2 million) and astrocytes (0.66 million) were seeded at a 3:1 ratio within each well in Astrocyte Media (0.5X B27 + 0.5X N2 + 1X GlutaMAX in DMEM/F12). Plates were centrifuged at 100g for 3 minutes until cells settled at the bottom. Half of the media were exchanged with fresh media every day. Microglia were seeded a week after initial Neuron-Astrocyte assembloid formation at one-third the number of astrocytes (0.2 million). Half the media was exchanged with iTF-Microglia media and doxycycline (2 µg/mL). Subsequently, half the media is then removed every other day and replaced with Astrocyte media supplemented with iTF-Microglia media cytokines.

## Tissue clearing for imaging cells expressing fluorescent proteins in iAssembloids and image processing

To obtain images of cells within the iAssembloids, hiPSC derived neurons expressing BFP, astrocytes expressing membrane GFP and microglia expressing mScarlet were seeded into iAssembloids with cells not expressing any fluorescent proteins at a 1:10 ratio. iAssembloids were then cleared with the ClearT2 protocol^77^. In brief, iAssembloids were first fixed with 4% paraformaldehyde (16% stock, diluted 1:4 in DPBS, Electron Microscopy Sciences, Cat. No. 15710) and then incubated in a 25% formamide (MilliporeSigma, Cat. No. F9037) /10% DPBS solution. Then, iAssembloids were incubated in a 50% formamide / 20% DPBS solution and imaged with a Leica confocal microscope. Images were then used for 3D projections (Video S1, related to Fig. 1) which were generated with Fiji^78^, using the 3D project module with the following settings: projection method: brightest point; axis of rotation: y-axis, slice spacing: 2, initial angle: 0,total rotation: 360, rotation angle increment: 10, lower transparency bound: 1, Upper transparency bound: 255, Opacity: 0, Surface depth cueing: 100, Interior depth cueing: 50, and interpolate on.

## Immunohistochemistry

iAssembloids were fixed with 4% paraformaldehyde in DPBS for 20 minutes on a shaker. iAssembloid were then washed with 0.33% Triton X-100 in DPBS 3 times for 5-10 minutes each time. Following washing, iAssembloids were blocked with 10% Normal Goat Serum + 1% BSA (blocking buffer) for an hour at room temperature. iAssembloids were incubated in antibodies diluted in blocking buffer overnight at 4°C. Following incubation, iAssembloids were washed with 0.33% Triton X-100 in DPBS 3 times for 5-10 minutes and incubated in secondary antibody diluted in blocking buffer for 1-2 hours at room temperature. iAssembloids were mounted and imaged on a confocal microscope. iAssembloids for NRF2 images were sectioned into 20 µm sections. Primary antibodies used for this study were as follows: rabbit anti-Iba1 (Wako, Cat. No. 019-19741), mouse anti-NeuN, clone A60 (MilliporeSigma, Cat. No. MAB377), mouse anti- S100 (β-Subunit) (MilliporeSigma, Cat. No. S2532), chicken anti-Tuj1 (Aves Labs, Cat. No. TUJ- 0020), rabbit anti-NRF2 (Abcam, Cat. No. ab62352, rabbit anti-cFOS (Abcam, ab214672).

## Nuclear isolation from iAssembloids for single-nuclei sequencing

Nuclear isolation was performed as previously described^60,79^. First, ∼300 iAssembloids were dounce homogenized in 5 ml of lysis buffer (0.25 M sucrose, 25 mM KCl, 5 mM MgCl_2_, 20 mM Tricine-KOH, pH 7.8, 1 mM DL-Dithiothreitol (DTT) (MilliporeSigma, Cat. No. D0632-1G), 0.15 mM Spermine tetrahydrochlorine (MilliporeSigma, Cat. No. S1141-1G), 0.5 mM Spermidine trihydrochloride (MilliporeSigma, Cat. No. S2501-1G), 1X protease inhibitor (MilliporeSigma, Cat. No. 4693159001), and RNAse Inhibitor (Promega, Cat. No. N2615) with 10 strokes.

Following initial dounce homogenization, IGEPAL CA-630 (MilliporeSigma, Cat. No. I8896- 50ML) was added to a final concentration of 0.3% and the sample was homogenized with 5 more strokes. Cells were then filtered through a 40 μm cell filter, and Optiprep was added to the cell solution for a final concentration of 25% Optiprep (MilliporeSigma, Cat. No. D1556-250ML). This solution was layered onto a 30%/40% Optiprep gradient and centrifuged at 10,000g for 18 minutes using a SW41-Ti rotor. The nuclei were collected at the 30%/40% Optiprep interface and quantified using the Countess FL Automated Cell Counter (ThermoFisher Scientific/Invitrogen, Cat. No. AMQAX2000). Droplet-based nuclear capture and library preparation were performed using the Chromium Single Cell Gene Expression workflow. Nuclei were brought to a concentration of 1000 nuclei/µL in 30% Optiprep solution before loading according to manufacturer’s specifications. 10,000 cells per sample were targeted for sequencing. Sample preparation and library generation were performed using the v3.1 10X 3’ single cell RNA sequencing library kit (10X Genomics, Cat. No.1000147) or the Fluent Biosciences PIPseq T2 3’ Single Cell RNA kit (FBS-SCR-T2-8-V4-1, FBS-SCR-T2-8-V3&V4-2, FBS-SCR-T2-8-V4-4. FBS-SCR-T2-8-V3&V4-6) cDNA fragment analysis was performed using the Agilent 4200 TapeStation System with the D5000 HS Kit (Agilent, Cat. Nos. 5067-5592, 5067-5593, 5067-5594). Sequencing parameters and quality control were performed as described by The Tabula Muris Consortium^80^.

## Multi-well Multielectrode Array (Axion)

iAssembloids 6 (Axion Biosystems, Cat. No. M384-tMEA-6B and 24 (Axion Biosystems, Cat. No. M384-tMEA-24W) well Cytoview MEA plates were coated with poly-L-ornithine (0.01%), fibronectin (1 µg/mL) and laminin (5 µg/mL) to facilitate iAssembloid adherence. iAssembloids were positioned into the well over the electrodes using a microscope. iAssembloids were equilibrated in BrainPhys™ Neuronal Medium (StemCell Technologies, Cat. No. 05790) 24 hours before plating to facilitate comparison to 2D monocultured neurons. Measurements were taken using the Maestro Edge multiwell microelectrode array (MEA) and Impedance system (Axion Biosystems) at 15 minute intervals with the Maestro MEA platform software (Axion Biosystems) with the “Neural” default setting.

### *2D* monocultured neurons

24 well Cytoview MEA plates (Axion Biosystems, Cat. No. M384-tMEA-24W) were coated with 0.1% PEI (MilliporeSigma, Cat. No. 03880) dissolved in 1X Borate Buffer (10mM Boric Acid, 2.5mM Sodium Tetraborate, 7.5mM NaCl, adjusted to pH 8.4) overnight at 37°C, washed with water, then dried at room temperature. Wells were then coated with BrainPhys™ media and 15 µg/mL Laminin. Cells were then seeded at a density of 200,000 cells per well on day 0 of differentiation. Measurements were taken using the Maestro Edge multiwell microelectrode array (MEA) and Impedance system (Axion Biosystems) at 3 x 5-minute intervals with the Maestro MEA platform software (Axion Biosystems) with the “Neural” default setting after 14 days in culture.

### CRISPRi-based pooled screens

The H1 CRISPRi library was packed into lentivirus as previously described^6^. Briefly, H1 plasmid library with the top 5 sgRNA per gene^27^ were transfected into HEK293s using the TransIT®- Lenti Transfection Reagent (Mirus, Cat. No. MIR 6606) along with third generation lentiviral packaging mix. Viral particles were harvested by removing the supernatant and filtering it through a 45 nm syringe filter. Virus was concentrated using cold Lentivirus Precipitation Solution (Alstem; Cat. No. VC100) was added to this filtered solution at a 1:4 ratio. The mixture was then centrifuged at 1500g for 30 minutes at 4°C. The supernatant was then removed, and the virus was resuspended in StemFlex. CRISPRi-NGN2 expressing iPSCs were then transduced at a MOI < 0.7. Cells expressing sgRNAs were then selected for with 2 μg/mL puromycin (Gibco/Thermo Fisher Scientific, Cat. No. A1113803) for 2 days as cells were expanded for screens. After expansion, cells were pre-differentiated and seeded into iAssembloids for screens. At days 1, 14 and 28 post-assembly of the iAssembloids, iAssembloid were harvested using a wide-orifice pipette tip and washed with DPBS, after which samples were frozen for later sample preparation. Genomic DNA was extracted with the NucleoSpin® Blood XL (Macherey Nagel, Cat. No. 740950.10) and samples were prepared for sequencing on an Illumina NextSeq 500 based on previously described protocols^81,82^

### Neurodegeneration library gene selection and cloning

sgRNAs targeting risk genes for neurodegenerative diseases were selected from GWAS and other published studies. Briefly, a manually curated list of genes associated with Alzheimer’s disease, Parkinson’s disease, ALS, Frontotemporal dementia, Huntington’s disease and Charcot-Marie-Tooth disease was assembled, drawing, among others, on previous publications ^83–87^ and the NCBI MedlinePlus database.

Protospacer sequences and gene targets are listed in Table S2. This small, focused library consists of 5 sgRNA targeting 376 genes. Pooled sgRNA libraries were generated using the same protocol as previously described^82,88^. Briefly, an oligonucleotide pool that encoded the library was synthesized by Agilent, amplified via PCR and then cloned into the pLG15 backbone^82^. HiPSCs expressing CRISPRi-NGN2 machinery were then transduced with the sgRNA library at an MOI < 0.3. Cells were then differentiated and utilized for screens in iAssembloids.

### Focused library secondary screens

146 sgRNAs, including 136 targeting sgRNAs (2 sgRNAs per gene) and 10 non-targeting sgRNAs were cloned into the pMK1334 CROP-seq vector^6^. sgRNA oligos were synthesized (top and bottom strands; IDT), annealed, and pooled in equal amounts (Table S3). hiPSCs expressing CRISPRi-NGN2 machinery were then transduced with the sgRNA library at an MOI < 0.3 and differentiated. Timepoints at 1 day after seeding into AggreWell™ 800 plates and 14 days after seeding into AggreWell™ 800 plates were taken. The following conditions were then tested: 2D monoculture, standard iAssembloids, 3D neurons and microglia, 3D neurons and astrocytes, and 3D neurons by themselves. Different media conditions were also tested including the normal condition (Standard w/cytokines), without supplementing any microglia media (Astrocyte w/cytokines), adding in microglia with microglia base media, but with no cytokines (Standard), adding in only astrocyte media with no cytokines (Astrocyte) and with BrainPhys™. Half the media was exchanged every day. Genes with consistent phenotypes across conditions were selected for CROP-seq.

### CROP-seq

Pooled sgRNA library (Table S3) consisting of 2 sgRNAs per targeted gene and 4 non-targeting controls was selected from focused library secondary screens. Libraries were cloned and transduced as described with the same method above in focused library secondary screen with an MOI < 0.30.

#### Sample preparation for CROP-seq

iAssembloids and 2D monocultured neurons were dissociated with the Papain (Worthington; Code: PAP2; Cat. No.LK003178) diluted in DMEM/F12 at 20U/mL Papain solution was warmed to 37°C and 500 μL was added to iAssembloids and incubated for 5 minutes. iAssembloids were triturated 15 times and filtered through a 22 μm cell filter. After filtering, neurons were selected for using the anti-PSA-NCAM microbeads kit (Miltenyi Biotec, Cat. No. 130-092-981), similarly to previously described methods^89^. Cells were centrifuged at 300g for 10 minutes and resuspended in 60 μL of PBS + 0.5% BSA (MilliporeSigma, Cat. No. A7979). To this suspension, 20 µL of Anti-PSA-NCAM microbeads were added. The mixture was incubated at 4°C for 15 minutes. Cells were spun down at 300g for 10 minutes and washed with PBS + 0.5% BSA. MACS was performed according to manufacturer’s specifications. After selection, purity was evaluated with flow cytometry. Sample preparation and library generation were performed using the v3.1 10X 3’ single cell RNA sequencing library kit. cDNA fragment analysis was performed using the Agilent 4200 TapeStation System. A portion of the cDNA was utilized to enrich for sgRNA sequences as previously described^6^. Sequencing parameters and quality control were performed as described by The Tabula Muris Consortium^80^.

### Flow cytometry

#### Cell Staining

Cells from iAssembloids and 2D neurons were dissociated with papain as described above and stained according to manufacturer’s protocols. Cells were washed three times with PBS after dissociation and incubated in DMEM/F12 with Liperfluo at 5 μM (Dojindo Molecular Technologies Inc., Cat. No. L24810) for 30 min or CellROX™ Orange Reagent at 2.5 μM (ThermoFisher Scientific/Invitrogen, Cat. No. C10443). Cells were then spun down and washed 2 more time with DPBS before analysis on the flow cytometer. Flow cytometry analysis was conducted with FlowJo v10.8.1.

#### GSK3_Β_ vs non-targeting guide survival assay

Neurons expressing two different *GSK3B* targeting guides and BFP were seeded at a 1:1 ratio with neurons expressing two different non-targeting guides expressing GFP. Seeding ratios were confirmed with flow cytometry before starting culture. After 14 days in culture in either iAssembloids or in monoculture, iAssembloids and neurons were dissociated with papain as described above and the number of cells expressing BFP (*GSK3B* knockdown) vs GFP (non-targeting guide) were quantified using FlowJo v10.8.1.

### Quantification of viability using trypan blue

Cells from iAssembloids and 2D neurons were dissociated with papain as described above and stained at 1:1 ratio with trypan blue solution (0.04%) (ThermoFisher Scientific/Gibco, Cat. No., 5250061). The cell and trypan blue mixture were then loaded into the Countess™ Cell counter. Two counts were taken for each sample and averaged. Average percent viability was recorded based on percent of trypan blue negative cells. Tetrodotoxin (Biotechne/Tocris, Cat. No. 1069) was used at a final concentration of 1.5 µM for a duration of 1 week, and ferrostatin 1 (Biotechne/Tocris, Cat. No. 5180) at 10 µM for a duration of 1 week.

### Western Blot

Approximately equal numbers of cells for both 2D monoculture, 3D monoculture and iAssembloids were lysed in 4X NuPAGE™ LDS Sample Buffer (ThermoFisher Scientific/Invitrogen, Cat. No., NP0007) + 10% 2-Mercaptoethanol, boiled for 10 minutes at 98 °C and then diluted 1:1 with DPBS. Samples were then loaded at equal volume into NuPAGE 4–12% Bis-Tris gels (ThermoFisher Scientific/Invitrogen, Cat. No. NP0336BOX). Gels were then transferred onto nitrocellulose membranes in Towbin buffer (25mM Tris base, 192 mM glycine, 20% (v/v) methanol) and blocked with 4% BSA in TBST (1X TBS + 0.1% Tween 20) followed by overnight incubation with primary antibodies at 4°C. Membranes were then washed 4X with TBST for 5 minutes and incubated with secondary antibodies at room temperature for 45 minutes. Membranes were imaged on the Odyssey Fc Imaging System (LI-COR, Cat. No. 2800) using Image Studio (v5.2) software. Primary antibodies used were mouse anti-GAPDH (Santa Cruz Biotechnology, Cat. No. sc-47724), rabbit anti-GSK3β (Cell Signaling, Cat. No. 12456S), and rabbit-anti S9 GSK3β (Cell Signaling, Cat. No. 9336S). Secondary antibodies were IRDye 680RD goat anti-mouse IgG (1:10,000 dilution, LI-COR, cat. no. 926-68070) and IRDye 800CW goat anti-rabbit IgG (1:10,000 dilution, LI-COR, cat. no. 926-32211).

### Data Analysis

#### snRNA sequencing for PIPseq and 10X Genomics

For 10X Genomics datasets, data processing was performed using CellRanger v4.0. Demultiplexing was performed using CellRanger mkfastq, a 10x-aware wrapper for bcl2fastq. Due to the large number of introns resulting from snRNA-seq, a pre-mRNA GTF was generated using the cellranger--mkref function. Briefly, GTF annotation rows were extracted based on the feature “transcript” of the original, pre-built GRCh38 GTF file. The feature types were then replaced from “transcript” to “exon.” Then cellranger--mkref was run to build a pre-mRNA GTF file. Alignment, filtering, barcode counting, and UMI counting was performed using cellranger count, though which gene expression matrices were generated. For PIP-seq datasets, alignment was performed using Fluent Bioscience’s software, PIPseeker, following options for snRNAseq. Outputs from cellranger and Pipseeker were then loaded into Seurat^90,91^, where nuclei with less than 500 UMI or greater than 15000 UMIs were filtered out. Doublets were then removed with DoubletFinder^92^. Using the NormalizeData() function in Seurat, raw counts per cell were normalized to the total expression in the cell, scaled to 10,000 transcripts per nucleus, and then log transformed. After normalization, 2000 of the most highly variable genes were identified using the FindVariableFeatures() function. The data were scaled such that the mean expression of each gene across all the cells is equal to 0 and the variance equal to 1, so that highly expressed genes do not dominate downstream analyses. Principal component analysis was performed using RunPCA() with default parameters and the first 10 PCAs were considered for clustering. Clustering was performed with the FindNeighbors() and FindClusters() functions at a resolution of 0.5. Uniform Manifold Approximation and Projection (UMAP) was calculated using the runUMAP() function in Seurat. Cell identities were then defined based on expression of cell type specific markers. Differentially expressed genes were identified by running FindMarkers() using the Wilcoxon-rank sum test.

Datasets from other publications^21,26^ were integrated using Seurat’s pipeline for mapping and annotating query datasets^90^. Briefly, data were preprocessed as above and a list of “anchor” genes between datasets were found using the function FindIntegrationAnchors(). The data is then integrated using the function IntegrateData().

#### Pathway enrichment analysis for snRNAseq

Pathway enrichment analysis for snRNAseq for neurons, astrocytes, and microglia were performed using EnrichR to classify assigned pathways for specific clusters^93–95^. Pathway analysis for snRNAseq for APOE3 or APOE4 astrocyte + neuron co-cultures were performed with The Database for Annotation, Visualization and Integrated Discovery (<u>DAVID</u>)^96,97^.

#### CRISPRi-based screens

Pooled screens were analyzed as previously described^6^, with the MAGeCK-iNC pipeline. First, sequencing reads were aligned to a reference file using Bowtie v0.99 to determine sgRNA frequencies per sample. Further analysis was then performed using the MAGeCK-inc software. Gene scores were calculated for each gene, which is defined as phenotype score multiplied by the -log10(P value). Hit genes were defined as genes with a false discovery rate < 0.05.

Volcano plots were generated with ggplot2 and heatmaps were generated with the pheatmap function in R.

#### Pathway enrichment analysis for CRISPRi-based screens

GO term enrichment analysis was performed using The Database for Annotation, Visualization and Integrated Discovery (<u>DAVID</u>)^96,97^ with the library gene list as the background list and hits from the screens as the genes of interest. *Reference sets include GOTERM_BP_DIRECT , BioCarta, BBID, KEGG_PATHWAY, REACTOME_PATHWAY and WIKIPATHWAYS were utilized.* Clustering was performed using DAVID and the enrichment score (mean of the -log(P) of annotation cluster) was calculated and groups were ranked based on the enrichment score.

#### Multiwell Multielectrode array (MEA) analysis

MEA analysis was performed with the default Axion’s Integrated Studio (AxIS) software. The default settings were used for spike detection with the Adaptive Threshold Crossing method. and network burst detection setting was applied. Spike files were then imported into the Axion neural metrics software where a 120 s snapshot of the total datafile was taken, and raster plots were generated.

#### CROP-seq differentially expressed genes analysis

sgRNA mapping was performed a previously published program and sgRNA assignment was performed using the program quantifyTranscriptome.R (https://github.com/powellgenomicslab/CROP-seq). Guides were only assigned if they were 1) the only guide found in the cell, 2) have more reads than three times the sum of other reads assigned to the other guides in the cell and the guides target a different gene, or 3) the guides for a single gene have 3x the sum of reads assigned to other guides in the cell. This information was then integrated as metadata for each sample as a Seurat object. UMAPs were generated with the same method as described above for snRNA-sequencing.

Differentially expressed genes were found using DESeq2 by pseudo bulking cells with the same sgRNA sequences by adapting previously described methods^98^. Cells with targeting guides versus non-targeting guides with individual sgRNA sequences counting as replicates. The P value cut off was 0.05. All differentially expressed genes were identified and compared log2FC for these genes within the cells with targeting guides. Differentially expressed genes with a log2FC magnitude greater than 2 in at least one condition (one gene target) was selected for heatmap visualization. Genes were manually annotated with some guidance from EnrichR. Differentially expressed genes for *GSK3B* was then independently examined and plotted using ggplot2.

#### Image analysis for NRF2 localization in iAssembloids

Three different iAssembloid sections for *GSK3B* KD and non-targeting control were taken and analyzed after staining with the protocol described above. Images were taken with a Leica SoRA confocal microscope. Nuclei were recognized as nuclear localized BFP+ (blue) objects while NRF2+ cells were recognized in a separate channel (red+). Nuclei were then masked onto the NRF2 stain to determine the location of the nuclei in relationship to NRF2 levels and then the intensity of the objects were then quantified using the MeasureObjectIntensity function.

## Data availability

snRNAseq (<u>GSE272094</u>), CROPseq (<u>GSE272093</u>) datasets are available on NCBI GEO. CRISPRi-based screens will be uploaded to CRISPRbrain (https://www.crisprbrain.org/). Protocols for iAssembloid culture, dissociation, and nuclear dissociation for snRNAseq will be uploaded to protocols.io.

## Supplemental Information Video, Tables and Legends

Video S1. 3D projection of BFP-labeled neurons sparsely seeded within iAssembloids

(related to Fig. 1).

3D projection of 36 images (Z stack; 1 um steps, 36 um total) taken of an iAssembloid with sparsely seeded BFP-labeled neurons (pseudocolored in white). iAssembloid was cleared with the ClearT2 method. Related to Figure 1.

Table S1. snRNA-seq: Differentially expressed genes from 2D monoculture vs neurons from iAssembloids and glial clustering information (related to Fig. 2)

The first tab provides differentially expressed genes from 2D monoculture neurons^20^ vs neurons from iAssembloids (Columns: differentially expressed genes, percent of cells with transcript found in monoculture population, percent of cells with transcript found in neurons of the iAssembloid population, and the adjusted P value). The second tab provides the Biological Processes GO terms from EnrichR. Columns: GO Terms (top 500 genes higher in log_2_FC in iAssemboids versus 2D monoculture), P value, adjusted P value, DEGs that fall within GO term, and the calculated -log10(adjusted P value). The third tab contains clustering information and number of astrocytes (columns: cluster number, number of cells in 4 week old iAssembloids, number of cells in 2 week old iAssembloids, number of cells in 1 day old iAssembloids, and in the monoculture conditions, as well as cluster assignments). The fourth tab consists of EnrichR terms from BioPlanet 2019 (columns: Term, overlap, p-value, adjusted p-value, odds ratio, combined scores, and genes). The fifth tab consists of EnrichR terms from GO Biological processes (columns: term, overlap, p-value, adjusted p-value, odds ratio, combined score, genes), and the final tab the same information for microglia (columns: cluster number, number of cells in 6TF microglia snRNAseq monoculture, microglia in the 2 week iAssembloid culture, microglia in the 4 week iAssembloid culture, relative percentages of each population, and annotated clusters).

**Table S2. Protospacer sequences for neurodegeneration library** (related to Fig. 3)

Protospacer sequences and sgRNA identities for neurodegeneration library used in primary screen. Columns: sgRNA identity (long format), protospacer sequence, gene target, and strand.

Table S3. Phenotypes from the primary screens using the H1 and neurodegeneration libraries and secondary survival screen in different media conditions sgRNAs and phenotypes (related to Fig. 3)

Phenotypes from the H1 library screen (day 14 iAssembloids, day 28 iAssembloids, day 14 neurons with TCW (serum) astrocytes, each in different tabs) and neurodegeneration library (day 14 iAssembloids, day 28 iAssembloids) are listed for all genes targeted H1 and neurodegeneration library, respectively (see Methods for details). Columns: targeted transcription start site, knockdown phenotype, P value, targeted gene, and the gene score (product of phenotype x –log_10_(P value)). The following tab provides protospacer sequences and phenotypes from the validation screens are listed for all sgRNAs from the secondary survival screen library (see Methods for details). For the protospacer sequences, columns are: sgRNA identifiers, protospacer sequences, gene target, and strand. The subsequent tabs provide phenotype information for the secondary validation screens (Media conditions: iAssembloids with astrocyte media, astrocyte media with cytokines, standard conditions, standard conditions with cytokines, BrainPhys™ Media; Culture conditions: 3D neurons only and 3D neurons with microglia). For the phenotypes from the different screens, the columns are: targeted transcription start site, knockdown phenotype, P value, targeted gene, and the gene score (product of phenotype x –log_10_(P value)). The final tab provides the output from DAVID GO platform for categorizing hits. Each cluster is annotated with the Enrichment score. Each column represents the category, the term, number of genes (count) that fit in the term, percent of the total, p-value, name of genes, list total, pop hits, pop total, fold enrichment, Bonferroni, Benjamini and FDR.

**Table S4. Differentially expressed genes from the CROP-Seq screen** (related to Fig. 4)

The first tab provides the numerical values underlying the heatmaps in Fig. 4B. Columns: genes targeted by CRISPRi. Rows: differentially expressed genes. The following tabs list changes in gene expression for *GSK3B* knockdown versus nontargeting sgRNAs in iAssembloids and in monoculture output from DESeq2. Columns: baseMean (average of the normalized count, divide by size factors over all cells), log2(baseMean) for the plot, log2FC, P value. NRF2 targets are annotated in the last column. See Methods for details.

Table S5. Phenotypes from primary H1 library screen in APOE-ε3 and APOE-ε4 astrocyte-neuron co-cultures (related to Fig. 7).

Phenotypes from the H1 library screen (day 14 KOLF2.1 APOE-ε3 Astrocytes cultured with KOLF2.1 APOE-ε3 neurons, day 14 KOLF2.1 APOE-ε4 Astrocytes cultured with KOLF2.1 APOE-ε3 neurons, day 14 TCW APOE-ε3 Astrocytes cultured with WTC11 APOE-ε3 neurons, and day 14 TCW APOE-ε4 Astrocytes cultured with WTC11 APOE-ε3 neurons) Columns: targeted transcription start site, knockdown phenotype, P value, targeted gene, and the gene score (product of phenotype x –log_10_(P value)).

Table S6. Differentially expressed genes from snRNAseq related in APOE-ε3 and APOE-ε4 astrocyte-neuron co-cultures (related to Fig. 7).

Tab 1 provides differentially expressed genes (DESeq2) from APOE-ε3 and APOE-ε4 astrocyte-neuron co-cultures. The first column are the gene names of the differentially expressed genes. Second column is the p-value, third column is the log2 fold change, third column is the percent represented by APOE-ε3 population, the third column represent percent expressing gene in the APOE-ε4 population, and the last column is the adjusted p-value. Tab 2 provides the Biological Processes GO terms from DAVID GO. Columns: GO Terms, Count: number of DEGs that fall into category, %: percent of terms in GO term, p-value, gene names represented in GO term, size of list in GO term, fold enrichment, and methods for multiple-hypothesis testing (Bonferroni and Benjamini-Hochberg), and false discovery rate. Last tab provides the output from Wilcoxon Rank-Sum test for unbiased clustering of excitatory neurons (RFBOX3+, SL) from Haney et al., 2024 dataset (Columns: gene name, p-value, average log2FC, pct.1, pct.2, adjusted p-value, cluster number)

---

**Figure S1.**
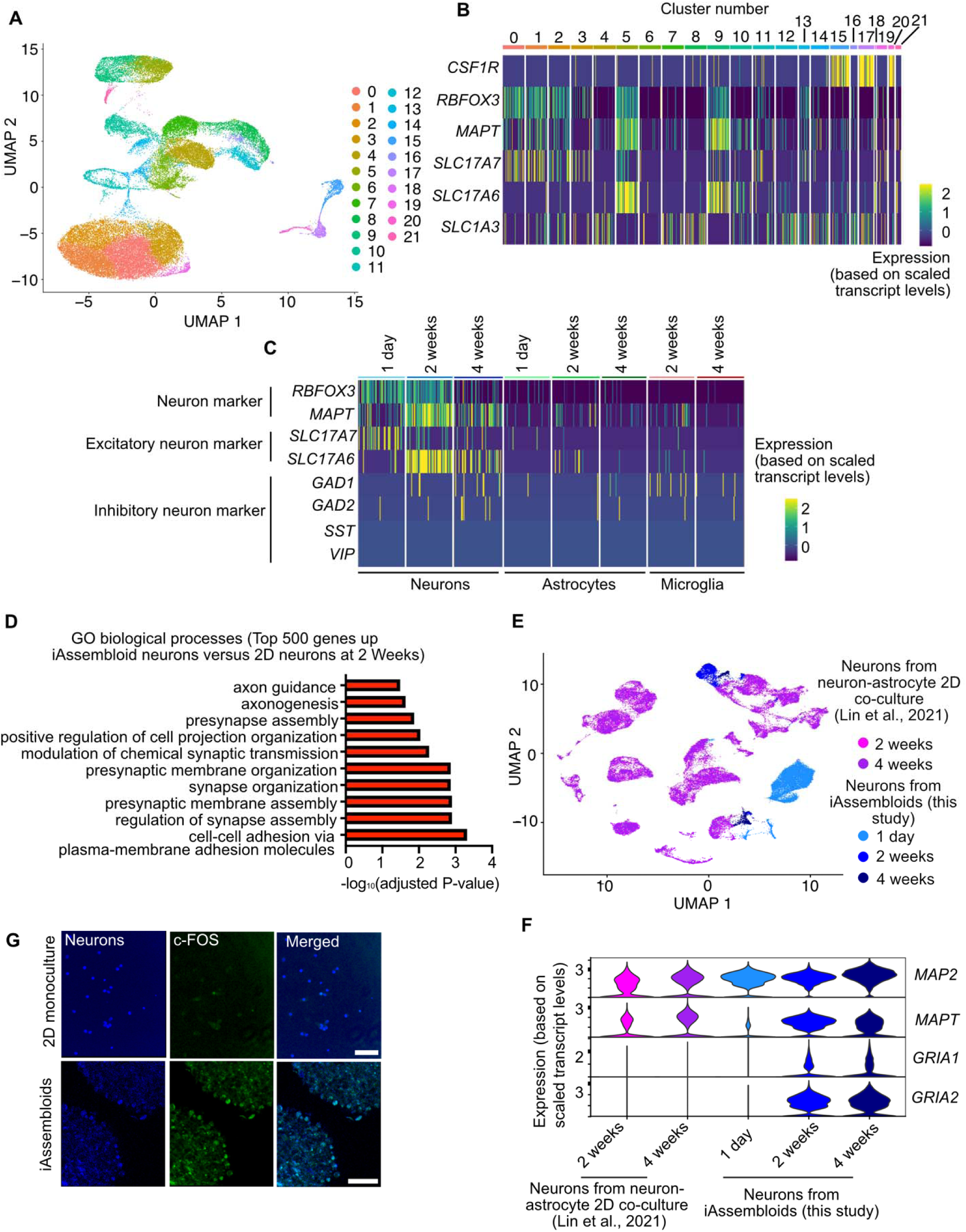
Single-nucleus RNA sequencing of iAssembloids (A) UMAP representation of snRNA sequencing data of iAssembloids. Colors represent clusters assigned through unbiased clustering (B) Heatmap representing expression of cell type-specific genes for cluster assignment. *CSF1R*: Microglia, *RBFOX3*, *MAPT*, *SLC17A7*, *SLC17A6*: Neurons, *SLC1A3*: Astrocytes (C) Heatmap representing expression of neuronal subtype-specific markers. *RBFOX3*, *MAPT*: Pan-neuronal markers, *SLC17A7*, *SLC17A6*: excitatory neuron markers, *GAD1*, *GAD2*, *SST*, *VIP*: inhibitory neuron markers. (D) Most significant Gene Ontology Biological Processes enriched in the 500 top genes expressed more highly in neurons in iAssembloids vs. 2D monocultured neurons. Adjusted P values were calculated using the EASE Score, a Modified Fisher Exact P-value and Benjamini-Hochberg method for correction for multiple hypothesis testing. (E) UMAP representation of integrated dataset from astrocyte and neuron 2D co-culture versus iAssembloids. (F) Selected differentially expressed genes between neurons from 2D astrocyte-neuron co-culture^22^ versus neurons from iAssembloids. (G) BFP+ neurons (blue) in monoculture and cultured within iAssembloids were stained for c-FOS (green). Scale bars represent 100 µm.

**Figure S2.**
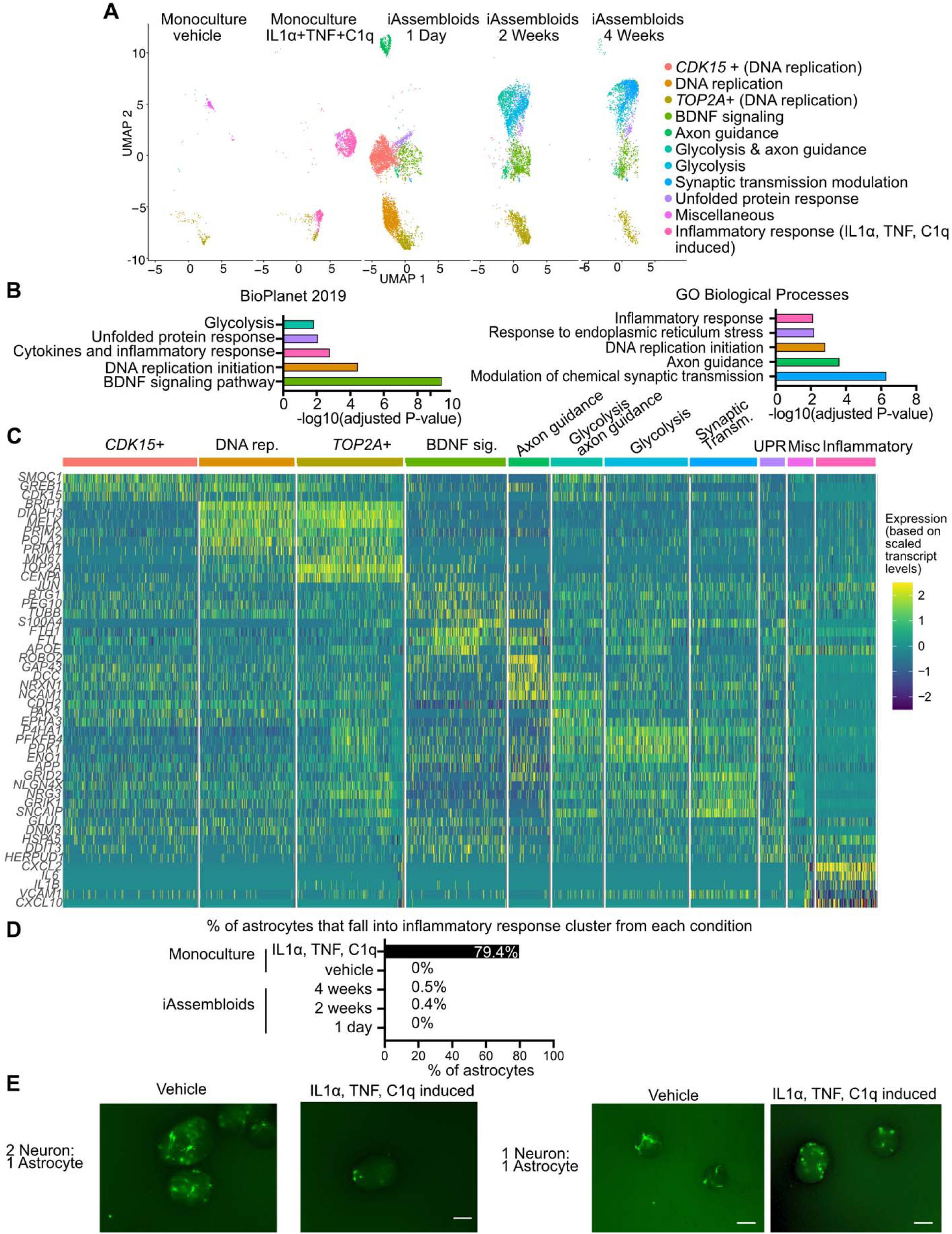
Astrocytes in iAssembloids express genes associated with neuronal support (related to Fig. 2) **(A)** Cells identified as astrocytes from snRNA-seq in iAssembloids (this study) were integrated with data from single cell sequencing of monocultured astrocytes^9^using Seurat’s data integration pipeline. A UMAP was generated and split by origin of sample and time point. Cluster names were defined by running defined markers through EnrichR. Clusters include a *CDK15*+ cluster, cells undergoing DNA replication, an immature astrocyte cluster *(TOP2A+)*, cells expressing genes related to BDNF signaling, axon guidance, glycolysis, synaptic transmission modulation, the unfolded protein response, and a cluster that had no specific markers (miscellaneous). The final cluster (IL1α, TNF, C1q induced) is based on previous studies for cytokine-induced markers^9^. **(B)** EnrichR BioPlanet 2019 pathway analysis was used to define clusters. Redundant terms (terms that consist of the same genes) were removed and only the term that had the highest - log_10_(adjusted p-value) is displayed. E.g. *PRIM2, POLA2, PRIM1* fall into the category of DNA replication initiation and Leading strand biosynthesis, but only DNA replication is displayed here. Colors represent clusters that fell into pathway/gene ontology categories.P represents Fisher’s Exact Test. Multiple correction testing is performed with the Benjamini-Hochberg method. **(C)** Heatmap was generated based on markers defined in A and B. **(D)** After clustering, the percent of total astrocytes that falls within the “inflammatory response category was plotted **(E)** Green channel represents astrocytes tagged with GFP sparsely seeded with neurons and non-fluorescent astrocytes. Astrocytes are seed at either a 2 neuron to 1 astroycte or 1 astrocyte to 1 astrocyte ratio. Representative images shown with iAssembloids treated with media (vehicle) and with media plus the triple cytokines (IL1α, TNF, C1q). The number of processes per astrocyte in focus (outlined in white) was manually counted. N=8 cells across 4 iAssembloids for vehicle and 8 cells across 3 iAssembloids for induced. Standard error of the mean is represented. P-value determined by student’s t-test. Scale bar represents 100 micron.

**Figure S3.**
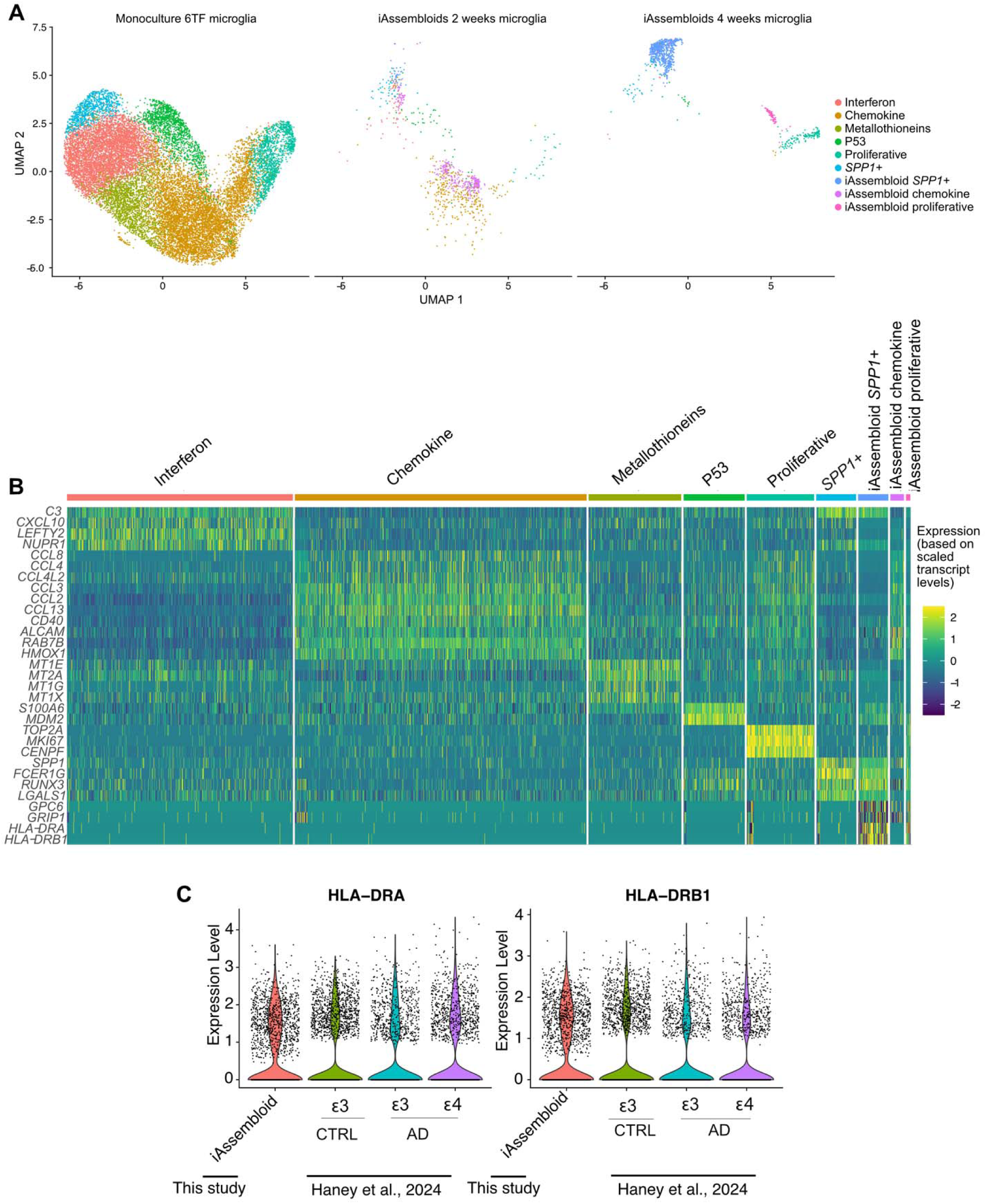
Microglia within iAssembloids express MHC Class II proteins. (related to Fig. 2) **(A)** Cells identified as microglia from snRNA-seq in iAssembloids were integrated with single-cell sequencing of microglia^8^ using Seurat’s data integration pipeline. A UMAP was generated and split by origin of sample and the time point. Clusters include those that fall into the interferon, chemokine, metallothioneins, P54, proliferative and *SPP1+* cluster. iAssembloid microglia specific clusters (iAssembloid *SPP1+,* iAssembloid chemokine, and iAssembloid proliferative are also highlighted). Microglia mainly map to existing clusters. However, cells in iAssembloids uniquely express MHC Class II proteins such as *HLA-DRA* and *HLA-DRB1*. **(B)** Heatmap was generated based on markers defined in A. **(C)** Expression level (log normalized value) of *HLA-DRA* and *HLA-DRB1* in a previously published patient dataset^26^ versus expression level of both genes from microglia grown in iAssembloids (this study)

**Figure S4.**
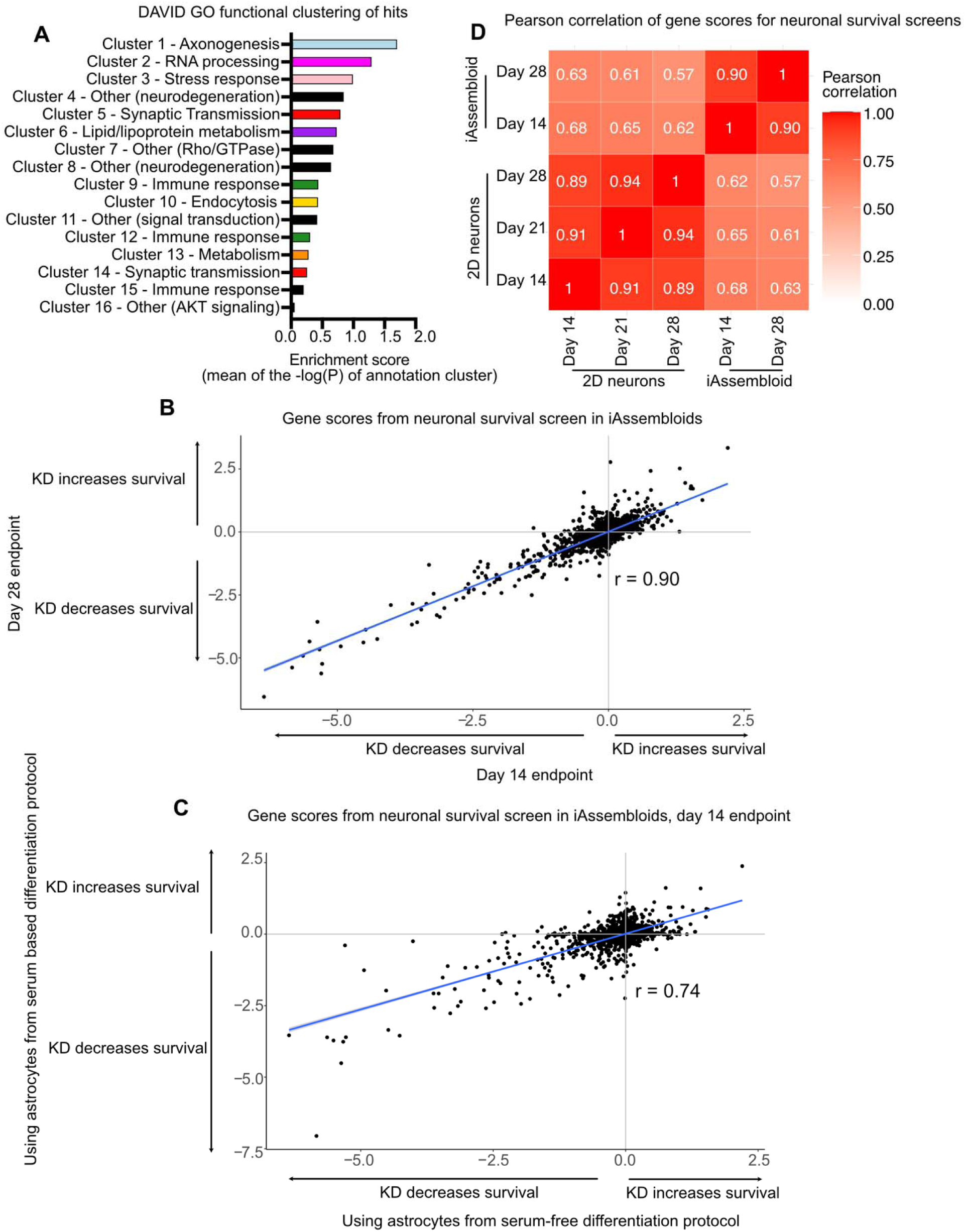
CRISPRi-based functional genomics screens in iAssembloids are reproducible (related to Fig. 3) **(A)** DAVID GO functional annotation clustering enrichment scores for 16 identified clusters. Enrichment score represents the mean -log_10_(P-value) of the terms within the annotation cluster. **(B)** Scatterplot comparing CRISPRi-based neuronal survival screens at 14 days vs. 28 days post seeding into AggreWell plates. **(C)** Scatterplot comparing CRISPRi-based neuronal survival screens using astrocytes from serum-based differentiation protocol^19,56^ vs. the serum-free differentiation protocol^13,14^. **(D)** Correlation heatmap of CRISPRi-based neuronal survival screens in the context of 2D monocultures^6^ vs. iAssembloids (this study).

**Figure S5.**
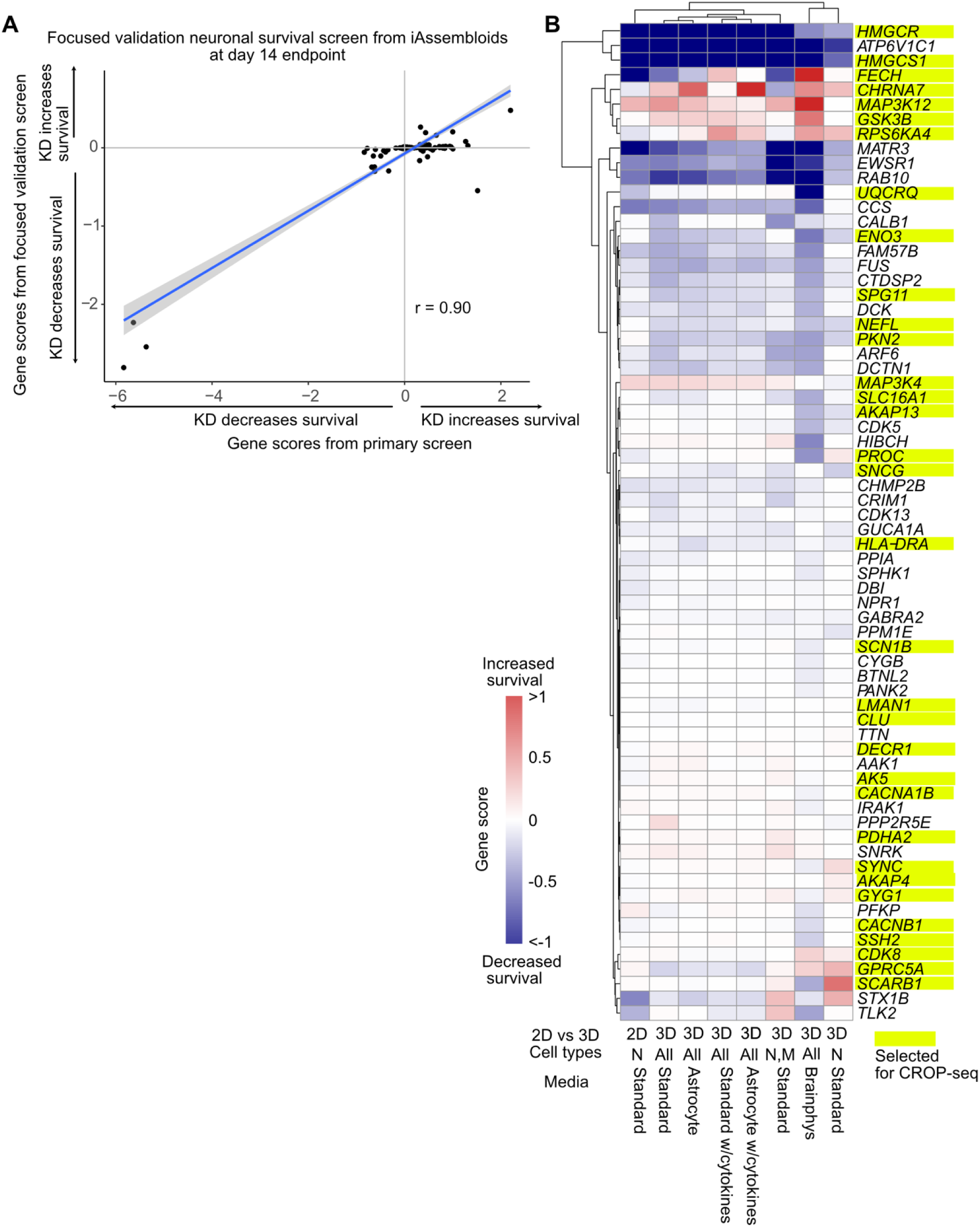
Focused validation screens for top hits from the primary screen. (related to Fig. 3 and 4) **(A)** Scatterplot of gene scores from the primary iAssembloid screen hits vs the secondary validation screen. Phenotypes from the secondary screen have a narrower dynamic range but are highly correlated with the primary screen (r = 0.9). **(B)** Results from secondary screens comparing iAssembloid culture in various media: normal media (composed of 0.5X astrocyte media and 0.5X microglia base media), normal media plus cytokines (our iAssembloid culture condition), astrocyte media, and astrocytes media plus cytokines, as well as 3D cultures with different cell type compositions: neurons only (N) and neurons plus microglia (N,M).

**Figure S6.**
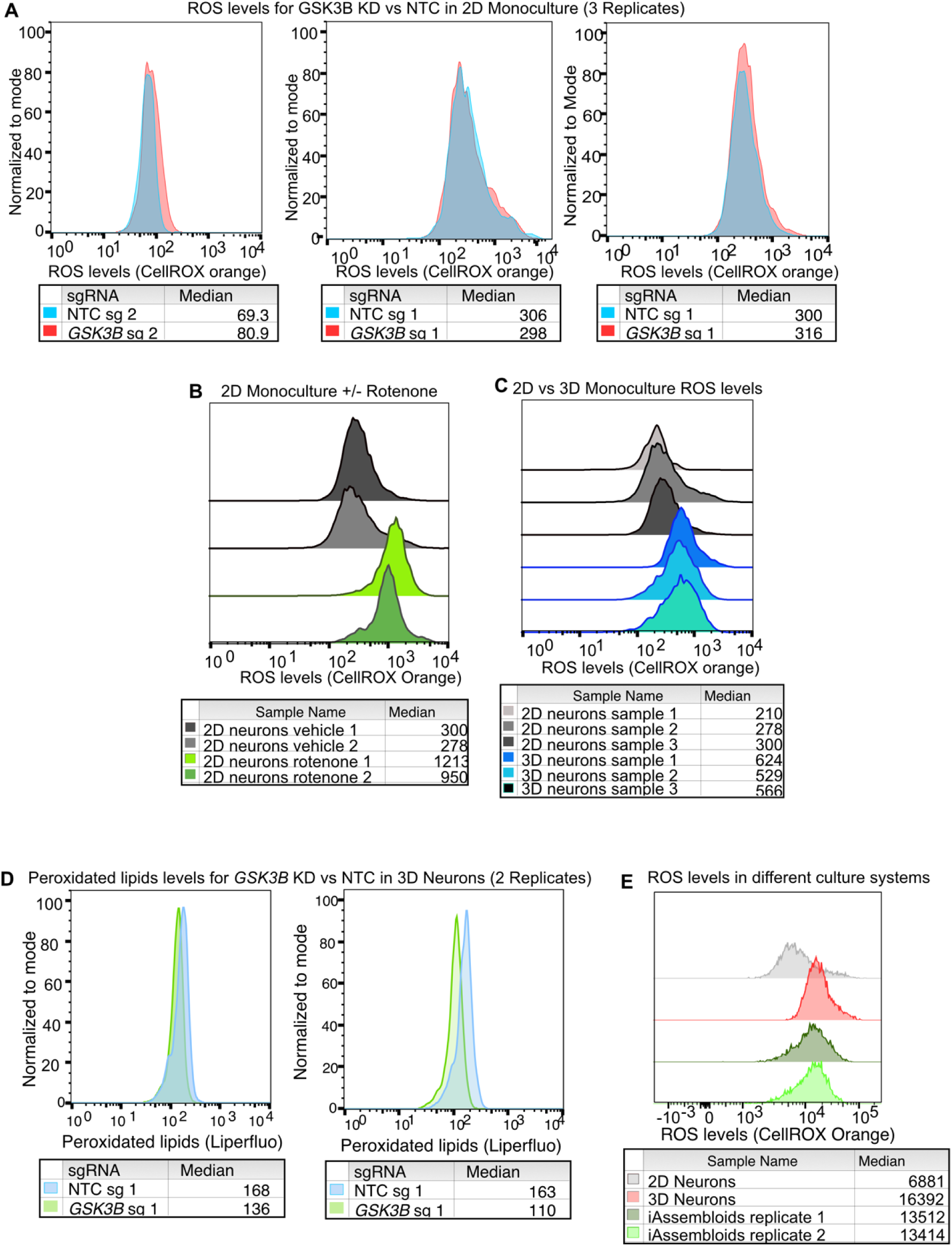
Representative flow cytometry results for neuronal phenotypes. (related to Fig. 5) **(A)** Three replicates of 2D monoculture neurons with and without *GSK3*Β knockdown stained with CellROX^TM^ orange. **(B)** Replicates of 2D vs 3D monocultured neurons stained for ROS levels by CellROX^TM^ orange. **(C)** Replicates of 2D monocultured neurons with and without rotenone treatment stained for ROS levels by CellROX^TM^ orange. **(D)** Replicates of 3D cultured neurons with and without GSK3Β knockdown stained with Liperfluo to detect peroxidated lipid levels **(E)** Neurons from 3 different culture systems (2D monoculture, 3D monoculture, iAssembloids) stained for ROS levels by CellROX^TM^ orange.

**Figure S7.**
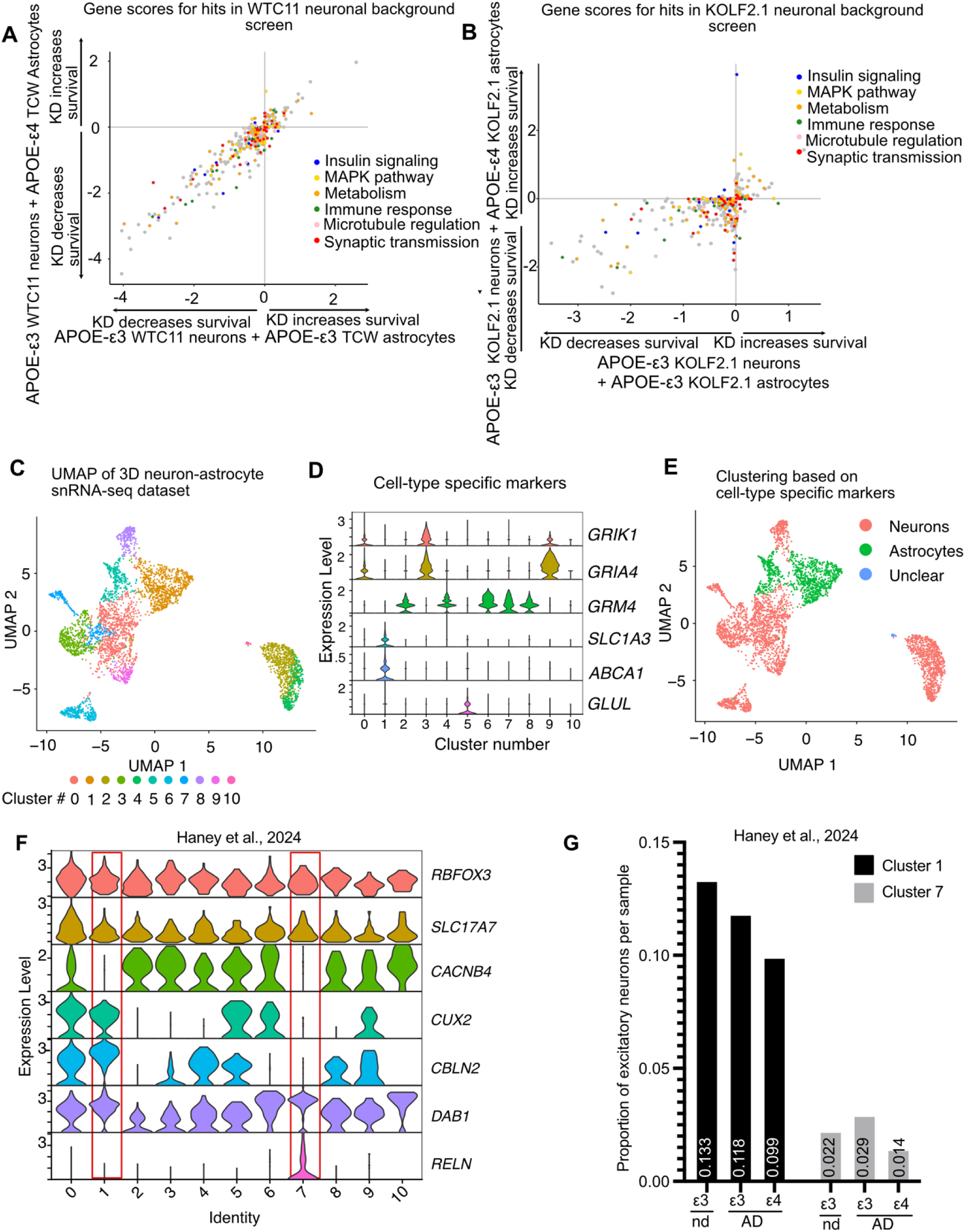
CRISPRi-based screen for neuronal survival in APOE-ε3 versus APOE-ε4 astrocyte 3D co-cultures and culture characterization (related to Fig. 7) **(A)** Scatterplot of gene scores from screens using APOE-ε3 versus (x-axis) compared to APOE-ε4 (y-axis) astrocytes (TCW 1E33-C, TCW 1E44-C) and APOE- ε3 neurons (WTC11). Hits (FDR < 0.05) from screens were included in the scatterplot. Genes belonging to selected functional categories were annotated in different colors; other genes are shown in gray. **(B)** Screen as described in B, but both neurons and astrocytes were generated in the KOLF2.1 background. **(C)** UMAP representation of snRNA sequencing data of 3D astrocyte (APOE-ε3 and APOE-ε4) and neuron (APOE-ε3) co-cultures. Colors represent clusters assigned through unbiased clustering **(D)** Cluster assignments based on expression of cell type-specific markers **(E)** UMAP of final assignments based on cell type-specific markers. Neurons are highlighted in red and astrocytes are highlighted in green. Cells that have unclear assignments were removed **(F)** Violin plot of excitatory neurons (RBFOX3+, *SLC17A1*+) cells identified from clusters previously generated by Haney et al., 2024. *CACNB4, CUX2, DAB1,* and *RELN* levels, which were shown to be differentially expressed by Wilcoxon Rank-Sum test were also plotted. **(G)** Proportion of neurons in clusters 1 and 7 in AD cases based on APOE status (ε3 vs ε4) compared to control (nd, ε3) neurons normalized to total number of excitatory neurons per sample.

## Notes

### Summary of Updates

We have conducted additional analyses to test our hypotheses in datasets from human brains, and clarified methodological aspects of our work.

